# The role of Tyr34 in proton-coupled electron transfer of human manganese superoxide dismutase

**DOI:** 10.1101/2024.05.29.596464

**Authors:** Jahaun Azadmanesh, Katelyn Slobodnik, Lucas R. Struble, Erika A. Cone, Medhanjali Dasgupta, William E. Lutz, Siddhartha Kumar, Amarnath Natarajan, Leighton Coates, Kevin L. Weiss, Dean A. A. Myles, Thomas Kroll, Gloria E. O. Borgstahl

## Abstract

Human manganese superoxide dismutase (MnSOD) plays a crucial role in controlling levels of reactive oxygen species (ROS) by converting superoxide (O_2_^•−^) to molecular oxygen (O_2_) and hydrogen peroxide (H_2_O_2_) with proton-coupled electron transfers (PCETs). The reactivity of human MnSOD is determined by the state of a key catalytic residue, Tyr34, that becomes post-translationally inactivated by nitration in various diseases associated with mitochondrial dysfunction. We previously reported that Tyr34 has an unusual pK_a_ due to its proximity to the Mn metal and undergoes cyclic deprotonation and protonation events to promote the electron transfers of MnSOD. To shed light on the role of Tyr34 MnSOD catalysis, we performed neutron diffraction, X-ray spectroscopy, and quantum chemistry calculations of Tyr34Phe MnSOD in various enzymatic states. The data identifies the contributions of Tyr34 in MnSOD activity that support mitochondrial function and presents a thorough characterization of how a single tyrosine modulates PCET catalysis.

## INTRODUCTION

About 25% of known enzymes perform redox reactions to promote cellular functions^1^. These enzymes, known as oxidoreductases, couple electron transfers with proton transfers in a process called proton-coupled electron transfer (PCET)^1–3^. PCETs are required for a wide range of biological functions, including energy generation and DNA synthesis, and oxidoreductase dysfunction leads to many disease states^4–8^. While PCETs are a fundamental biochemical reaction, how they are facilitated by enzymes is not well understood due to difficulties in unraveling the precise proton transfer and electron transfer steps^2,9,10^. Some enzymes use multiple proton transfers per electron transfer step or vice-versa^11–13^. Defining how PCET reactions occur in oxidoreductases benefits the understanding of redox-mediated disease states and furthers therapeutic interventions^14–18^.

Some oxidoreductases use their PCETs to regulate the concentrations of reactive oxygen species (ROS) in cells. These PCETs are crucial as fluctuations of ROS levels stimulate mitophagy and programmed cell death, and excessive levels lead to the damage of DNA, proteins, and lipids^19^. Impairment of oxidoreductase function promotes several diseases, including cardiovascular and neurological conditions and cancer progression^20,21^. In particular, the oxidoreductase human manganese superoxide dismutase (MnSOD) is responsible for eliminating O_2_^•−^ in the mitochondrial matrix, and dysfunction of its activity contributes to a broad range of diseases^22^.

Within the mitochondrial matrix, MnSOD uses PCETs to decrease O_2_^•−^ concentrations rapidly and efficiently (*k*_cat_/K_m_ > ∼ 10^9^ M^-1^ s^-1^). In the first half-reaction, O_2_^•−^ is oxidized to molecular oxygen (O_2_) with a Mn^3+^ ion (*k*_1_), and in the second half-reaction, O_2_^•−^ is reduced to hydrogen peroxide (H_2_O_2_) with a Mn^2+^ ion (*k*_2_). The two half-reactions regenerate the Mn^3+^ ion^23,24^. O_2_^•−^ is endogenously produced as a byproduct of the mitochondrial electron transport chain, and MnSOD is the only means to reduce O_2_^•−^ in the mitochondrial matrix^25,26^. Aberrations of MnSOD function play roles in psoriasis, inflammatory bowel disease, multiple sclerosis, cardiovascular disease, and breast and prostate cancer^27–31^. Overall, the PCETs of MnSOD contribute to health and longevity^24,32^.

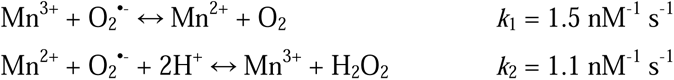

For PCET catalysis, MnSOD uses a hydrogen bond network (dashed blues lines, **Fig. 1a**) to couple proton transfers with changes in Mn oxidation state^32–34^. The Mn ion is covalently bound to His26, His74, His163, Asp159, and a solvent molecule called WAT1. From WAT1, the hydrogen bond network extends to Asp159, Gln143, Tyr34, another solvent molecule denoted as WAT2, His30, and Tyr166 from the adjacent subunit. To enter the active site, the substrate and solvent pass through a gateway formed by Tyr34 and His30. Hydrophobic residues Trp123, Trp161, and Phe66 pack the hydrogen bonding atoms of Asp159, WAT1, Gln143, and Tyr34 close together^35^. In our previous work, we investigated the protonation states of the hydrogen bond network in relation to the oxidation state of the Mn ion^13^.

**Fig. 1:**
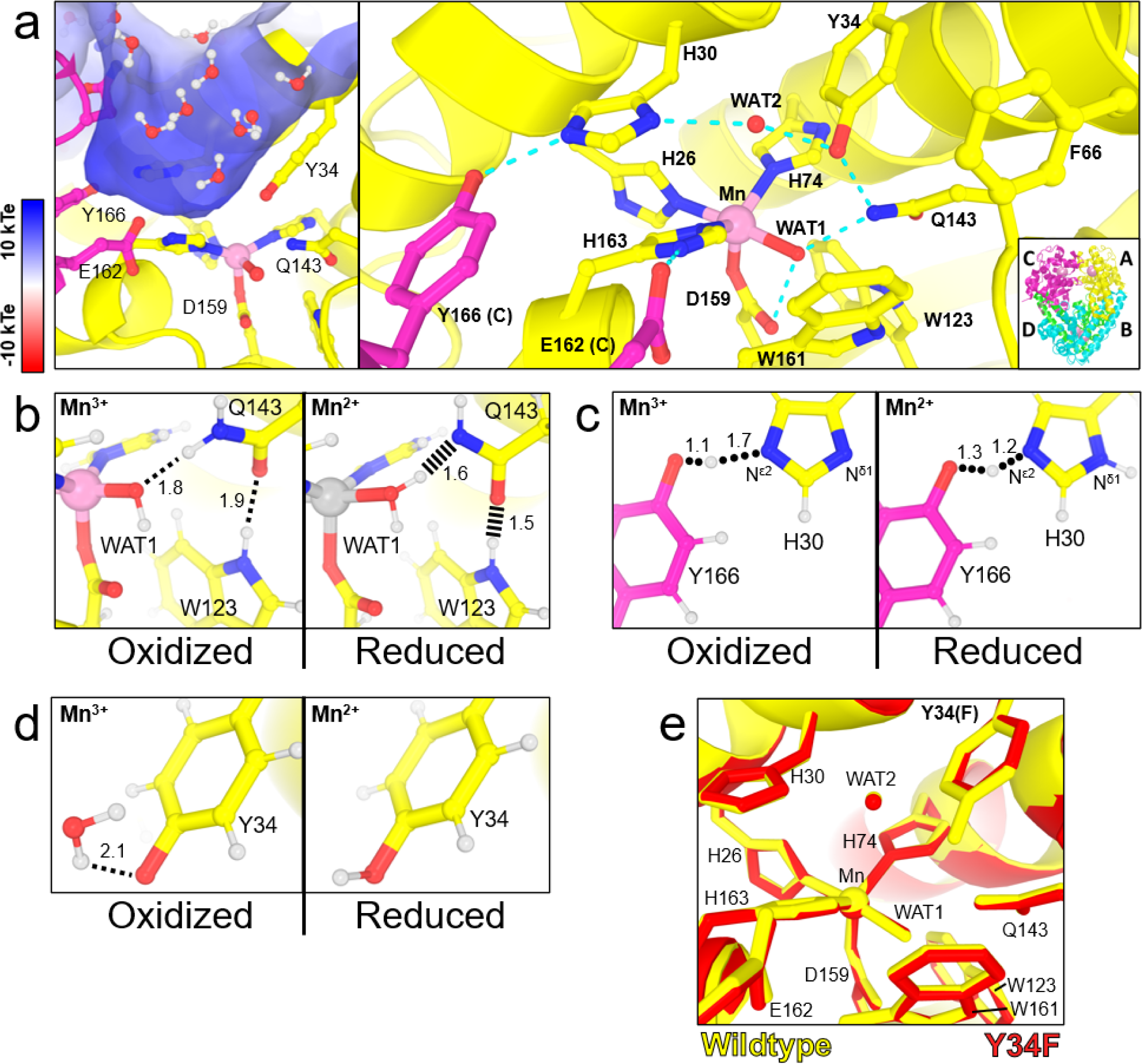
Structure of human wildtype MnSOD and protonation differences between oxidation states. **a** The active sites of tetrameric MnSOD are found in positively charged cavities made by two subunits. Solvent and substrate must pass through gateway residues His30 and Tyr34 to interact with the catalytic Mn ion. Blue dashed lines indicate hydrogen bonds. The inset indicates the chain identity derived crystallographically, where the asymmetric unit is composed of an AB dimer, and the CD dimer is generated through symmetry to form the native tetrameric assembly. **b** Room temperature neutron structures revealed changes in protonation among WAT1 and Gln143 and alterations in the hydrogen bond network of WAT1-Gln143-Trp123. Dotted lines indicate hydrogen bonds ≥ 1.8 Å while hashed lines indicate SSHBs that are hydrogen bonds < 1.8 Å. **c** A shared proton, indicated with rounded dots, was observed between His30 and Tyr166. This proton helps modulate the protonation state of N^δ1^(His30) when the Mn ion changes oxidation states. **d** Tyr34 is deprotonated in Mn^3+^SOD and protonated in Mn^2+^SOD. **e** Superposition of wildtype MnSOD (yellow, PDB ID 5VF9) and Tyr34Phe MnSOD (red, PDB ID 9BWR) active sites with a root-mean-square deviation of 0.07 Å among C^α^ atoms. Panel **a** was created from MnSOD X-ray structure (PDB ID 5VF9)^71^, and panels **b-d** were created from the neutron structures of Mn^3+^SOD (PDB ID 7KKS) and Mn^2+^SOD (PDB ID 7KKW)^13^. All hydrogen positions were experimentally determined except for solvent molecules in panel **a** that were randomly generated to accentuate the solvent in the active site funnel. All distances are in Å.

Using neutron crystallography, we previously observed how changing the Mn oxidation state shifts the pK_a_ of active site amino acids and leads to several unusual protonation states^13^. Neutron diffraction is advantageous for studying PCET mechanisms because the scattering of deuterium is on par with carbon, nitrogen, and oxygen, and neutrons do not introduce photoreduction of metal ions, unlike X-rays^36,37^. The neutron structures of Mn^3+^SOD and Mn^2+^SOD revealed three important pieces of data on how PCET catalysis is facilitated. First, during the Mn^3+^ to Mn^2+^ redox transition, an unconventional proton transfer occurs between Gln143 and metal-bound ^−^OH(WAT1) leading to an unusual Gln143 amide anion and H_2_O (**Fig. 1b**). The amide anion is stabilized by two short-strong hydrogen bonds (SSHBs) with WAT1 and Trp123. SSHBs stabilize catalytic steps and enhance kinetic rates (hashed lines, **Fig 1b**)^38–40^. Second, a shared proton between Tyr166 and N^ε2^(His30) modulates the protonation of N^δ1^(His30), with N^δ1^(His30) only being protonated in the Mn^2+^ oxidation state (**Fig. 1c**). A low-barrier hydrogen bond (LBHB) is formed between Tyr166 and His30 in the Mn^2+^ state. A LBHB is a type of SSHB where the heteroatoms transiently share a proton, and the hydrogen bond distances between the donor D/H atom and the heteroatoms are nearly equivalent (1.2-1.3 Å)^41^. Third, Tyr34 is deprotonated in the Mn^3+^ oxidation state and becomes protonated in the Mn^2+^ state. Tyr34 and N^δ1^(His30) are probably the two proton donors needed during the Mn^2+^ to Mn^3+^ redox transition where H_2_O_2_ forms from the protonation of the substrate (*k*_2_)^32,34,42,43^. Overall, the neutron structures indicate that multiple proton transfers are coupled to electron transfer events and that the active site metal leads to residues with unconventional pK_a_s that are essential for PCET catalysis.

How O_2_^•−^ interacts with the active site for catalysis is unclear^32–34^. O_2_^•−^ may bind the Mn ion directly for an “inner-sphere” electron transfer or between His30 and Tyr34 for a long-range “outer-sphere” electron transfer with the Mn ion^32–34,43^. Quantum chemistry computational studies have postulated that the first half-reaction (*k*_1_, Mn^3+^ → Mn^2+^) proceeds through an inner-sphere electron transfer while the second half-reaction (*k*_2_, Mn^2+^ → Mn^3+^) proceeds through an outer-sphere electron transfer^43^. The outer-sphere mechanism for the second half-reaction is a promising hypothesis because His30 and Tyr34 have been shown to lose protons during the Mn^2+^ → Mn^3+^ redox transition that converts O_2_^•−^ to H_2_O_2_ (**Fig. 1c, d**)^13^. His30 and Tyr34 could protonate O_2_^•−^ in concert with long-range electron transfer to form the H_2_O_2_ product. While there is no experimental evidence for how O_2_^•−^ interacts with the active site for electron transfer, modeling and quantum chemical calculations support the second half-reaction proceeding through an outer-sphere mechanism^43^.

Human MnSOD is inhibited by its product, H_2_O_2_, to regulate the output of mitochondrial H_2_O_2_. Physiologically, MnSOD product inhibition is thought to be related to H_2_O_2_ acting as a secondary messenger^33,34,44^. Mitochondrially-derived H_2_O_2_ plays a role in apoptosis^45,46^, mitochondrial biogenesis^44^, and protein localization and activity^47^. Furthermore, mitochondrial H_2_O_2_ has been shown to modulate the activity of phosphatase and tension homolog (PTEN), a protein tyrosine phosphatase that contributes to cancer and neurological disease^48,49^. While mitochondrial H_2_O_2_ may be scavenged by peroxiredoxins (PRDXs) and glutathione peroxidases (GPXs), these enzymes are dependent on nicotinamide adenine dinucleotide phosphate (NADPH) for function^50–52^. MnSOD product inhibition regulates mitochondrial H_2_O_2_ levels independent of metabolic status. However, little is known about how MnSOD mechanistically achieves product inhibition.

Product inhibition occurs when the ratio of O_2_^•−^ to MnSOD is high^33,34^. At concentrations of O_2_^•−^ that are lower than enzyme, catalysis proceeds through the first-order reactions *k*_1_ and *k*_2_. At concentrations of O_2_^•−^ that are much greater than enzyme, catalysis proceeds through the reversible inhibition reactions *k*_3_ (first-order) and *k*_4_ (zero-order). The *k*_3_ kinetics are attributed to forming a “product-inhibited” complex that is unreactive to O_2_^•−^ and only disassociates through the zero-order *k*_4_ reaction that dictates the lifetime of the complex. Several kinetic models suggest that inhibition is initiated from the reaction of a Mn^2+^ ion with O_2_^•−^ in the absence of protonation to form [Mn^3+^-O_2_^2−^] or [Mn^3+^-^−^OOH] (*k*_3_)^33,34^. The inhibited complex is then relieved by at least one protonation to yield Mn^3+^ and H_2_O_2_ (*k*_4_). Since *k*_2_ and *k*_3_ both use Mn^2+^ and O_2_^•−^ as reactants, they are competing reactions, and their ratios determine the propensity for MnSOD to become product-inhibited. Human MnSOD has equal *k*_2_ and *k*_3_ (**Table 1**), which means at high O_2_^•−^ to enzyme ratios, ∼50% of Mn^2+^ reactions with O_2_^•−^ form the inhibited complex^53,54^. The factors that govern *k*_3_ and *k*_4_ are unclear though the residue Tyr34 appears to contribute to the inhibition process^55,56^.

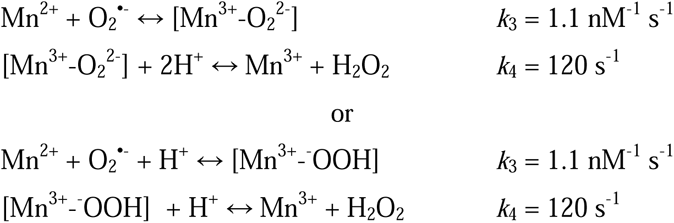

**Table 1.** Individual steady-state rate constants of MnSOD.

| **MnSOD** | $k_1$ ($\text{nM}^{-1} \text{s}^{-1}$) | $k_2$ ($\text{nM}^{-1} \text{s}^{-1}$) | $k_3$ ($\text{nM}^{-1} \text{s}^{-1}$) | $k_4$ ($\text{s}^{-1}$) |
| --- | --- | --- | --- | --- |
| Human Wildtype<sup>a</sup> | 1.5 | 1.1 | 1.1 | 120 |
| Human Y34F<sup>b</sup> | 0.55 | <0.02 | 0.46 | 52 |
| Human W161F<sup>a</sup> | 0.30 | <0.01 | 0.46 | 33 |
| Human W123F<sup>b</sup> | 0.76 | <0.02 | 0.64 | 79 |

<sup>a</sup>pH 8.2 from Hearn *et al.*<sup>35</sup>
<sup>b</sup>pH 7.5 from Greenleaf *et al.*<sup>55</sup>

Tyr34 is a conserved residue in the active site of MnSOD^55,57^. Physiologically, the residue becomes nitrated in several human neurodegenerative diseases, leading to inactivated MnSOD^58–65^. Mechanistically, Tyr34 is thought to protonate dioxygen species to produce H_2_O_2_ during the Mn^2+^ to Mn^3+^ redox transition^32,34,57^. This is in line with our previous neutron structures, where Tyr34 is poised to donate a proton in Mn^2+^SOD (**Fig. 1d**)^13^. With mutation of the residue to phenylalanine, Tyr34Phe MnSOD is unable to proceed through the fast Mn^2+^ to Mn^3+^ redox transition (*k*_2_), and catalysis proceeds exclusively through the product-inhibited pathway (*k*_3_ **Table 1**). Since *k*_2_ << *k*_3_, ∼99% of Mn^2+^ reactions with O_2_^•−^ form the inhibited complex. There is also higher retention of the inhibited complex, with half the disassociation rate compared to wildtype (*k*_4_, **Table 1**). The perturbed kinetics and enrichment of product inhibition from the Tyr34Phe variant could be rationalized by the loss of a hydroxyl group for proton transfer, especially since the active site of the Tyr34Phe variant from the X-ray structure is nearly identical to that of the wildtype (**Fig. 1e)**^57^. Interestingly, the Trp161Phe variant, which does not eliminate a hydroxyl group, has similar kinetics to the Tyr34Phe variant (**Table 1**)^35^. This suggests that the contribution of Tyr34 toward catalysis is not solely proton transfer. To study the effects that govern the reactions *k*_1_-*k*_4_ of MnSOD, we performed a series of experiments with the Tyr34Phe variant.

Several studies have reported that mixing large H_2_O_2_ concentrations with Mn^3+^SOD leads to the formation of the product-inhibited complex that decays to Mn^2+^SOD^35,66–68^. Mutagenesis studies found the process to be correlated with the forward reactions *k*_2_-*k*_4_. These puzzling results may be explained by excessive concentrations of H_2_O_2_ initiating a backward reaction with Mn^3+^SOD to produce O_2_^•−^, and then a subsequent forward reaction of O_2_^•−^ with MnSOD to form the inhibited complex^67,68^. Importantly, this would provide an avenue to structurally analyze product inhibition without relying on volatile O_2_^•−^ solutions^69^. We combined H_2_O_2_ soaking with the Tyr34Phe variant to study the product-inhibited complex.

Here, we sought to elucidate the role of Tyr34 in MnSOD reactivity by utilizing the Tyr34Phe variant in combination with neutron crystallography, X-ray absorption spectroscopy (XAS), and quantum mechanical (QM) chemistry calculations. With neutron crystallography, we captured the product-bound, reduced, and oxidized states of Tyr34Phe MnSOD free from radiation-induced perturbations. For each state, we used XAS to probe the electron orbitals of the Mn ion. Then, we used QM calculations to quantitatively interrogate the Mn ion orbitals that determine redox activity. Lastly, the Tyr34Phe data were compared with wildtype^13^ and Trp161Phe^70^ to piece together a MnSOD mechanism that describes the reactions *k*_1_-*k*_4_. With respect to oxidoreductases in general, our work presents a thorough characterization of how a single tyrosine modulates PCET catalysis.

## RESULTS AND DISCUSSION

### Capture of MnSOD product inhibition with H_2_O_2_ soaking

With XAS, we sought to determine whether product inhibition of MnSOD may be achieved with excess H_2_O_2_. In Mn K-edge XAS, the X-ray absorption near edge structure (XANES) contains information on the oxidation and coordination states, while the extended X-ray absorption fine structure (EXAFS) region provides information on Mn-ligand bond distances. For studying product inhibition, the Tyr34Phe MnSOD variant is advantageous because it accumulates and retains the product-inhibited complex (**Table 1**)^33,34,55,57,67,72^. First, we pursued the XANES spectral signatures of Tyr34Phe Mn^3+^SOD and Tyr34Phe Mn^2+^SOD to establish reference points of oxidation and coordination state. Then, we compared the XANES spectra for Tyr34Phe Mn^3+^SOD mixed with O_2_^•−^ or H_2_O_2_.

The energy of the rising edge of the XANES region increases with higher oxidation states while the intensity of the pre-edge contains information about the coordination of the Mn ion. The pre-edge corresponds to 1s to 3d orbital transitions, with gains in intensity from 4p character mixing into the 3d orbitals from symmetry distortions and/or loss of inversion symmetry^73–75^. Based on electron paramagnetic resonance (EPR) data, the Mn ion is expected to be high-spin in both Mn^3+^ and Mn^2+^ states^76^. For oxidized Tyr34Phe MnSOD, the rising edge is observed at higher energy compared to the reduced data, indicating a difference in the Mn ion valency. Given that these samples were mixed with stoichiometric excesses of redox reagents, we interpret oxidized Tyr34Phe MnSOD as Mn^3+^ and reduced Tyr34Phe MnSOD as Mn^2+^. For both oxidized and reduced Tyr34Phe MnSOD, the pre-edge intensity (i.e., the area under the curve, inset **Fig. 2a**) corresponds to a distorted five-coordinate trigonal bipyramidal complex that is also observed in the crystal structures^57,77^. Orbital assignment to the pre-edge intensities will be addressed later with the Kα high-energy resolution fluorescence detected absorption (HERFD) method and excited state simulations. The spectral signatures of Tyr34Phe Mn^3+^SOD and Tyr34Phe Mn^2+^SOD were measured for comparison with the product-inhibited complex.

**Fig. 2:**
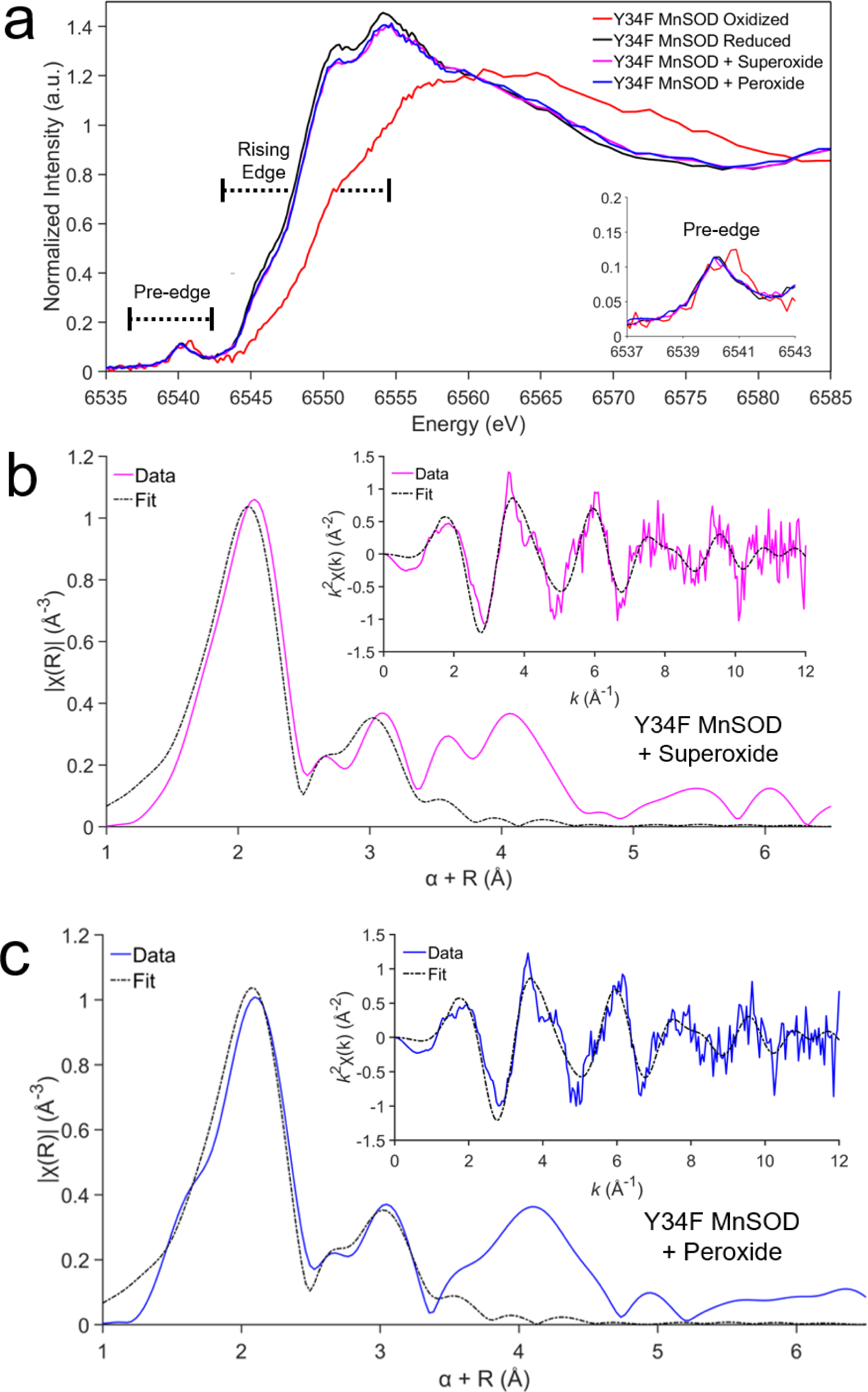
X-ray Absorption Spectroscopy of Tyr34Phe MnSOD. **a** The XANES region of Tyr34Phe MnSOD treated with potassium dichromate to isolate the Mn^3+^SOD resting state, sodium dithionite to isolate the Mn^2+^SOD resting state, and either superoxide or hydrogen peroxide to isolate the product-inhibited state. **b-c** Fourier transform of Mn K-edge EXAFS data [*k*^2^χ(*k*)] from superoxide-soaked and peroxide-soaked Tyr34Phe MnSOD with the EXAFs spectrum in *k*-space shown in the inset. Due to the scattering phase shift, the distance found by the Fourier Transformation (R) is ∼0.5 Å shorter than the actual distance and a 0.5 Å correction (α) was implemented. Colored lines represent experimental data, while the black line is simulated EXAFS spectra fit to the experimental data.

To isolate the product-inhibited complex with XAS, we either introduced O_2_^•−^ or H_2_O_2_ to Tyr34Phe MnSOD. O_2_^•−^ was used to isolate the complex from reaction *k*_3_ (**Table 1**), while H_2_O_2_ was used to attain the complex with a back reaction^67,78^. Following the methods of previous studies, the superoxide-soaked inhibited complex was attained by mixing O_2_^•−^ dissolved in dimethyl sulfoxide (DMSO) and di-benzo-18-crown-6-ether with Tyr34Phe MnSOD dissolved in aqueous solution^66,67,79,80^. The peroxide-soaked inhibited complex was obtained by simply mixing aqueous H_2_O_2_ with aqueous MnSOD. For both approaches, the inhibited complex has been reported to form within 44 ms of mixing with Tyr34Phe MnSOD^67^. Indeed, the approaches yield nearly identical XANES spectra and reveal that the same electronic structure is formed (**Fig. 2a**). The superoxide and peroxide-soaked spectra are most similar to Tyr34Phe Mn^2+^SOD, suggesting the presence of a divalent Mn ion. However, these spectra have unique features relative to Tyr34Phe Mn^2+^SOD, with lower intensities at ∼6550-6555 eV and higher intensities at ∼6565-6575 eV. Likewise, the pre-edge intensities are highly similar to Tyr34Phe Mn^2+^SOD to indicate that the resulting complex from either mixing O_2_^•−^ or H_2_O_2_ is five-coordinate (**Fig. 2a**). Altogether, by using the Tyr34Phe variant that enriches for the product-inhibited complex, we show that an electronically distinct five-coordinate Mn^2+^ complex forms from either exposure to O_2_^•−^ or H_2_O_2_.

Next, we investigated the EXAFS region of the superoxide and peroxide-soaked Tyr34Phe MnSOD K-edge spectra. The Fourier transform of the EXAFS data, χ(*k*), yields χ(*R*) that provides information on the atomic radial distribution around the absorbing Mn ion (**Fig. 2b, c**). Overall, the spectra in both *k* space and *R* space are highly similar between the soaked Tyr34Phe MnSOD counterparts and yielded the same Mn bond distance solutions (**Table 2**). For both superoxide and peroxide-soaked samples, the first shell of coordination observed at ∼2.1 Å is best fit by three N atoms at 2.15 Å and two O atoms at 2.11 Å to indicate a five-coordinate complex (**Supplementary Table 1, 2**). These account for N^ε2^ atoms of His26, His74, and His163, O^δ2^ atom of Asp159, and another O atom (denoted O^1^). As the next closest scatterer is a single atom found at 2.44 Å (denoted O^2^), we interpret O^1^ and O^2^ as belonging to a dioxygen species. The largest contributors to the second peak seen at ∼3.0 Å are seven carbon atoms that correspond to the C^δ2^ and C^ε1^ of the three histidines and the C^γ^ of the aspartate residue. Overall, the EXAFS data suggests that upon introducing O_2_^•−^ or H_2_O_2_ to Tyr34Phe MnSOD, a five-coordinate Mn complex that includes a dioxygen species is formed.

**Table 2.** Comparison of superoxide and peroxide-soaked Tyr34Phe MnSOD bond distances found through EXAFS fitting, neutron crystallography, and DFT calculations.

| Exp. | EXAFS (Å) | EXAFS (Å) | Neutron Structure (Å) |
| --- | --- | --- | --- |
|  | Y34F MnSOD + Superoxide | Y34F MnSOD + Peroxide | Y34F MnSOD + Peroxide |
| Mn-N<sup>ε2</sup>(H26) | 2.15 | 2.15 | 2.17 |
| Mn-N<sup>ε2</sup>(H74) | 2.15 | 2.15 | 2.23 |
| Mn-N<sup>ε2</sup>(H163) | 2.15 | 2.15 | 2.21 |
| Mn-O<sup>δ2</sup>(D159) | 2.11 | 2.11 | 2.13 |
| Mn-O<sup>1</sup>(LIG) | 2.11 | 2.11 | 2.00 |
| Mn-O<sup>2</sup>(LIG)<sup>a</sup> | 2.44 | 2.44 | 2.34 |
| Calc. | DFT (Å) | DFT (Å) | DFT (Å)<sup>b</sup> |
|  | Y34F Mn<sup>2+</sup>SOD + <sup>-</sup>O<sub>2</sub>H | Y34F Mn<sup>3+</sup>SOD + <sup>-</sup>O<sub>2</sub>H | Y34F Mn<sup>2+</sup>SOD + <sup>•</sup>O<sub>2</sub>H |
| *S* | 5/2 | 2 | 3 |
| Mn-N<sup>ε2</sup>(H26) | 2.28 | 2.00 | 2.12 |
| Mn-N<sup>ε2</sup>(H74) | 2.20 | 2.07 | 2.16 |
| Mn-N<sup>ε2</sup>(H163) | 2.16 | 2.19 | 2.17 |
| Mn-O<sup>δ2</sup>(D159) | 2.17 | 2.01 | 2.01 |
| Mn-O<sup>1</sup>(LIG) | 2.10 | 1.82 | 2.37 |
| Mn-O<sup>2</sup>(LIG)<sup>a</sup> | 2.38 | 2.60 | 3.03 |

<sup>a</sup>The second oxygen of the dioxygen ligand is not directly bound to the Mn ion.
<sup>b</sup>Broken-symmetry geometry optimization of Mn<sup>2+</sup>-<sup>•</sup>O<sub>2</sub>H with *S* = 2 collapses to a Mn<sup>3+</sup>-<sup>-</sup>O<sub>2</sub>H complex.

Our XAS data indicates that H_2_O_2_ has both a reducing effect on MnSOD and can form the inhibited complex. The reducing effect of MnSOD by H_2_O_2_ has been reported previously^66,67^. Thermodynamically, the process is unfavorable as the reduction potential of wildtype Mn^3+^SOD to Mn^2+^SOD is 0.4 V, and the oxidation potential of H_2_O_2_ to O_2_^•−^ is -0.85 V^81^. However, a 100-fold excess of H_2_O_2_ was used and probably drove a backward reaction^67,68^. Mutagenesis studies have indicated that forming the inhibited complex through peroxide-soaked is dependent on reactions *k*_2_-*k*_4_ and suggests that the O_2_^•−^ generated from the backward reaction then reacts with MnSOD in the forward direction^67^. In these past experiments, detection of O_2_^•−^ was lost during the dead time of the instrument (1.4 ms). As H_2_O_2_ is more stable than O_2_^•−^ in aqueous solution, we chose to use H_2_O_2_ for further structural studies of product inhibition.

### Neutron structure of the Tyr34Phe product-inhibited complex

To visualize the product-inhibited complex and the corresponding active site protonation states, we solved a cryocooled neutron structure of perdeuterated Tyr34Phe MnSOD soaked with deuterium peroxide (D_2_O_2_) at 2.30 Å resolution. Neutron crystallography is advantageous because it probes H/D atom positions and is absent of photoreduction effects that metal-oxygen species interactions are susceptible to from X-ray exposure^37^. For the neutron structures, proton positions can be assigned at 2.5 Å or better. We first solve the all-atom structure of the entire enzyme, excluding the active site. Then, with these phases, we carefully interrogate the active site nuclear density maps for the active site coordination and protonation states.

For D_2_O_2_-soaked Tyr34Phe MnSOD, we first analyzed the omit |*F*_o_| - |*F*_c_| nuclear scattering-length density of the active sites for any visually distinct dioxygen species. For chain A, nuclear density next to the Mn ion is observed at 3.0 σ and is oblong (**Fig. 3a**). We interpreted the density as a dioxygen species with a single proton (denoted as LIG for ligand) that has taken the place of the WAT1 position observed in the resting states (**Fig. 1b**). The proton of LIG, D^2^, points toward Trp161 and appears to be interacting with the pi system (**Fig. 3a**). The O^2^(LIG) and D^ε21^(Gln143) atoms are close in proximity (2.2 Å apart) though LIG and Gln143 are not in optimal geometry for hydrogen bonding (**Fig. 3a, b**). LIG refined well at full occupancy and led to Mn bond distances that closely resemble those measured from the EXAFS spectra (**Table 2**). At physiological temperatures, optical absorption spectra also suggest a displacement of WAT1 and binding of a dioxygen species, which agrees with our cryocooled diffraction data^82^. For chain B, the omit |*F*_o_| - |*F*_c_| is instead a spherical shape at 3.0 σ (**Supplementary Fig. 1b**), indicating a ^-^OD molecule that has been previously observed in wildtype neutron structures^13^. Contrasting structures between the two subunits of the asymmetric unit are often observed for the MnSOD *P*6_1_22 crystal form due to differences in solvent accessibility^13,71^. In chain B, the nuclear density near the Mn ion and ^−^OD molecule is difficult to interpret and will not be discussed further (**Supplementary Fig. 1b**). Regardless, from the nuclear density at chain A, a singly protonated dioxygen species replaces WAT1 upon D_2_O_2_ soaking and forms a complex with bond distances similar to those found from EXAFS spectra (**Table 2**).

**Fig. 3:**
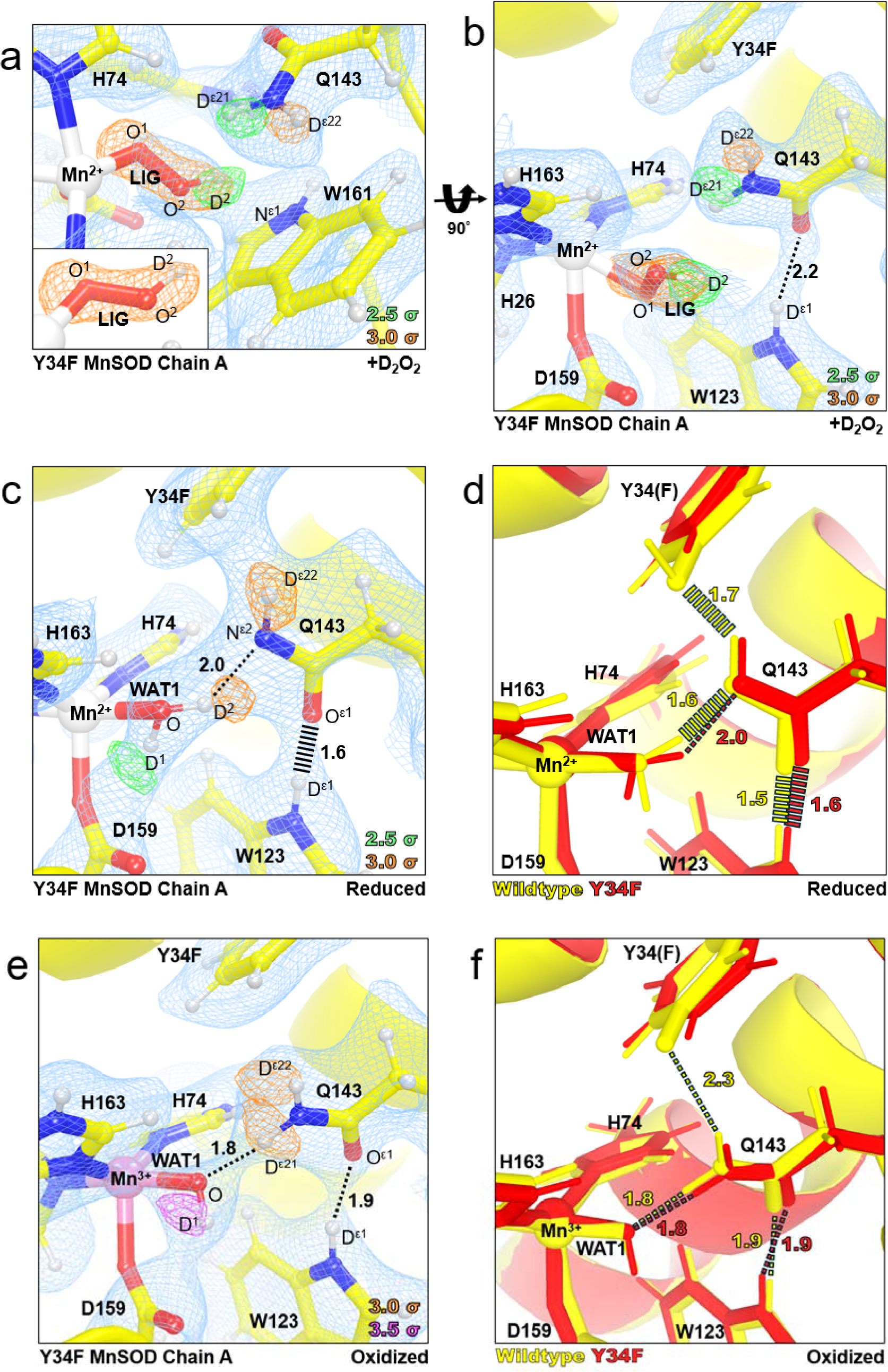
Neutron structures and protonation states at the active site of D_2_O_2_-soaked, reduced, and oxidized Tyr34Phe MnSOD. **a** D_2_O_2_-soaked Tyr34Phe MnSOD for chain A with a singly-protonated dioxygen species denoted LIG. The inset highlights the elongated |*F*_o_| - |*F*_c_| difference density of LIG. **b** D_2_O_2_-soaked Tyr34Phe MnSOD for chain A rotated 90° along the horizontal axis relative to panel **a**. **c** Tyr34Phe Mn^2+^SOD for chain A. **d** Active site overlay of wildtype Mn^2+^SOD (PDB 7KKW, solved at 2.3 Å resolution) and Tyr34Phe Mn^2+^SOD (this study, solved at 2.5 Å resolution). **e** Tyr34Phe Mn^3+^SOD for chain A. **f** Active site overlay of wildtype Mn^3+^SOD (PDB 7KKS, solved at 2.3 Å resolution) and Tyr34Phe Mn^3+^SOD (this study, solved at 2.3 Å resolution). Green, orange, and magenta omit |*F*_o_| - |*F*_c_| difference neutron scattering length density of protons displayed at 2.5σ, 3.0σ, and 3.5σ. respectively. Light blue 2|*F*_o_| - |*F*_c_| density is displayed at 1.0σ. Distances are in Å. Dashed lines indicate hydrogen bonds ≥ 1.8 Å, and hashed lines indicate SSHBs that are hydrogen bonds < 1.8 Å. Chain B active sites are shown in **Supplementary Fig. 1**.

How the product-inhibited complex is formed as well as its oxidation state and coordination, have been historically controversial^33,34,68,83–85^. The generally accepted mechanism is that inhibition is initiated from O_2_^•−^ binding Mn^2+^ to form a Mn^3+^-dioxo complex (*k*_3_, **Table 1**)^32–34,86,87^. Our XAS data indicates a five-coordinate Mn^2+^ as the identity of the complex^66,67^. Indeed, the neutron structure of Tyr34Phe MnSOD soaked with D_2_O_2_ yields a five-coordinated active site with a singly-protonated dioxygen species. The Mn bond distances of the neutron structure resemble those found from the XAS data and density function theory (DFT) calculations using a hydroperoxyl anion, ^−^O_2_H (**Table 2**). Overall, our data verifies that the inhibited complex is a five-coordinate Mn^2+^ complex where the WAT1 position has been replaced with ^−^O_2_H.

### Tyr34 orients the Gln143-WAT1 SSHB of Mn^2+^SOD and limits product inhibition

We next wondered whether enrichment of the product-inhibited complex in the Tyr34Phe variant was because of structural perturbations near the Mn ion or due to the absence of a hydroxyl group for proton transfer. To this end, we solved neutron structures of reduced and oxidized Tyr34Phe MnSOD at 2.50 and 2.30 Å resolution, respectively. Redox reagents were used that enacted their chemical effects without entering the active site^13,88^. For reduced Tyr34Phe MnSOD, the Mn bond distances are similar to five-coordinate wildtype Mn^2+^SOD (**Supplementary Table 3**). Furthermore, the protonation states resemble those of wildtype Mn^2+^SOD, where WAT1 is of the D_2_O form while Gln143 is deprotonated to the amide anion (**Fig. 3c**). Deprotonated amino acids are identified when attempts to model and refine a proton result in negative |*F*_o_| - |*F*_c_| difference neutron scattering length density and all the other protons of the amino acid can be placed. Interestingly, the lack of the hydroxyl group in Tyr34Phe Mn^2+^SOD slightly perturbs the orientation of Gln143 and lengthens the WAT1-Gln143 hydrogen bond (**Fig. 3d**). The WAT1-Gln143 SSHB of the wildtype enzyme is critical for the back-and-forth proton transfers needed for redox cycling of the Mn ion^13^, and the Tyr34Phe Mn^2+^SOD neutron structure suggests one role Tyr34 plays in catalysis is correctly positioning Gln143 for rapid PCET catalysis. Indeed, the Mn^2+^ to Mn^3+^ redox transition is nearly ablated for Tyr34Phe MnSOD (*k*_2_, **Table 1**)^33,34^. Another consequence of the perturbation of the WAT1-Gln143 interaction is the potentially easier displacement of WAT1 by a dioxygen species to form the product-inhibited complex. Overall, the reduced Tyr34Phe MnSOD neutron structure suggests that Tyr34 orients Gln143 for a tight hydrogen bonding interaction with WAT1.

For oxidized Tyr34Phe MnSOD, the Mn ion is bound by an ^−^OD molecule (**Fig. 3e**) and has covalent bond distances that resemble that of wildtype Mn^3+^SOD (**Supplementary Table 3**). Furthermore, the hydrogen bond distances among the interactions of O(WAT1)-D^ε21^(Gln143) and D^ε1^(Trp123)-O^ε1^(Gln143) are identical to wildtype counterpart (**Fig. 3f**). Of potential consequence for the Tyr34Phe variant, however, is the absence of the anionic phenolate group that is seen in wildtype Mn^3+^SOD. As PCET mechanisms depend on the distribution of electrostatic vectors to propagate proton and electron transfers, the lack of an ionizable group may perturb enzyme kinetics. Such electronic effects are suggested by the lower kinetic rates of the Mn^3+^ to Mn^2+^ redox transition found for the Tyr34Phe variant (*k*_1_, **Table 1**). Overall, the interactions between WAT1, Gln143, and Trp123 are the same in Tyr34Phe and wildtype Mn^3+^SOD.

The neutron structures of reduced and oxidized Tyr34Phe Mn^2+^SOD suggest the roles of Tyr34 in catalysis are (1) orient Gln143 for efficient interaction and proton transfer with WAT1 during the Mn^2+^ to Mn^3+^ PCET reaction, (2) limit formation of the product-inhibited complex by strengthening the WAT1-Gln143 hydrogen bond, and (3) contribute subtle electronic effects as a phenolate anion during the Mn^3+^ to Mn^2+^ reaction. These interpretations are supported by the kinetic rates of Tyr34Phe MnSOD^33,34^. For Tyr34Phe MnSOD, the fast Mn^2+^ to Mn^3+^ redox reaction is ablated (*k*_2_, **Table 1**), formation of the product-inhibited complex is enriched (*k*_3_ >> *k*_2_, **Table 1**), and the Mn^3+^ to Mn^2+^ redox reaction is cut in third (*k*_1_, **Table 1**). A unifying theme among Tyr34Phe MnSOD and other variants studied kinetically is that the precise orientation of Gln143 is critical for catalysis. Mutating residues directly adjacent to Gln143, such as Trp161, Trp123, and Tyr34, lead to similar kinetic consequences, where the Mn^2+^ → Mn^3+^ half reaction is ablated, product inhibition is enriched, and the Mn^3+^ → Mn^2+^ half reaction is slower compared to wildtype^33,34^. These redundant effects of the Trp161Phe, Trp123Phe, and Tyr34Phe variants suggest that a crucial role of residues neighboring Gln143 is to enforce a SSHB between WAT1 and Gln143. In the case of Tyr34, our neutron structures indicate that Tyr34 is responsible for positioning Gln143 for a SSHB with WAT1 during the Mn^2+^ resting state and that the strength of the bond between Gln143 and WAT1 correlates with the extent of product inhibition.

### Retention of the product-inhibited complex is dependent on the Gln143 position

As the XAS and neutron crystallographic data of Tyr34Phe MnSOD demonstrated the formation of a five-coordinated Mn^2+^SOD with a singly-protonated dioxygen species replacing the WAT1 position, we wondered if the formation and retention of the inhibited complex could be distinguished with other variants of MnSOD. In the context of Tyr34Phe MnSOD, we also sought to define whether the enrichment of product inhibition was due to the loss of the ionizable Tyr group only or also due to the perturbation of the Gln143- WAT1 SSHB interaction. For example, the Trp161Phe MnSOD variant alters a residue directly adjacent to Gln143 and has enriched product-inhibition kinetics like those of Tyr34Phe MnSOD (**Table 1**)^66^. For wildtype, Tyr34Phe, and Trp161Phe, we performed XANES in the HERFD mode of detection, which allows a large improvement in energy resolution and sensitivity compared to conventional XANES^89–91^. Here, we focused on comparing the reduced form of the variants to those of peroxide-soaked counterparts. As all these complexes are expected to be five-coordinated d^5^ with spin S=5/2, we sought to distinguish fine details of the spectra offered by the HERFD mode of detection.

For HERFD-XANES data of Tyr34Phe MnSOD, the features of the reduced and peroxide-soaked forms seen in conventional XANES (**Fig. 2a**) are also observed. While both forms have similar rising edge energies to indicate the same oxidation state of the Mn ion, the peroxide-soaked form is seen with a lower intensity at ∼6550-6555 eV and a higher intensity at ∼6565-6580 eV (**Fig. 4a**). Unique to the HERFD data is the observation of a higher intensity shoulder of peroxide-soaked Tyr34Phe MnSOD at ∼6565 eV and the ability to better resolve the pre-edge peaks for both samples. The peak centers of the pre-edge are within ∼0.2 eV of each other, while greater intensity for the higher energy tail (∼6542.5 eV) is distinguishable for peroxide-soaked Tyr34Phe MnSOD. Overall, reduced and peroxide-soaked Tyr34Phe MnSOD spectra have distinct features between each other, and the general shape trends are reproducible both with conventional XANES (**Fig. 2a**) and HERFD-XANES (**Fig. 4a**) and suggest two differing five-coordinate Mn^2+^ complexes are measured.

**Fig. 4:**
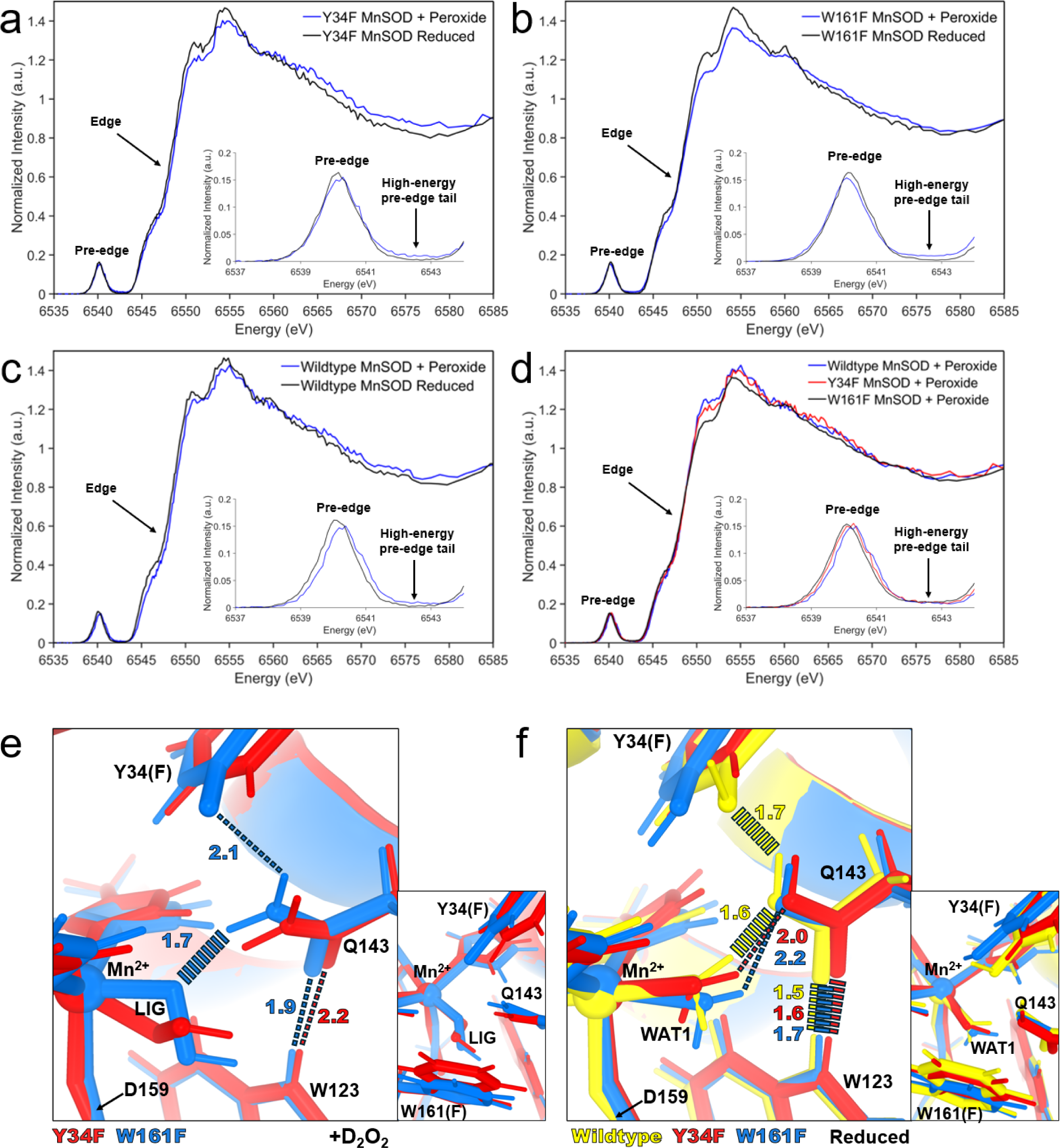
HERFD-XANES and neutron structures of Tyr34Phe, Trp161Phe, and wildtype MnSOD. **a** HERFD-XANES spectra of H_2_O_2_-soaked and reduced Tyr34Phe MnSOD. **b** HERFD-XANES spectra of H_2_O_2_-soaked and reduced Trp161Phe MnSOD. **c** HERFD-XANES spectra of H_2_O_2_-soaked and reduced wildtype MnSOD. **d** HERFD-XANES spectra of H_2_O_2_-soaked wildtype MnSOD, H_2_O_2_-soaked Tyr34Phe MnSOD, and H_2_O_2_-soaked Trp161Phe MnSOD. **e** Active site overlay of D_2_O_2_-soaked Tyr34Phe MnSOD (this study) and D_2_O_2_-soaked Trp161Phe MnSOD (PDB ID 8VHW, solved at 2.3 Å resolution). **f** Active site overlay of wildtype Mn^2+^SOD (PDB ID 7KKW, solved at 2.3 Å resolution), Tyr34Phe Mn^2+^SOD (this study, solved at 2.5 Å resolution), and Trp161Phe Mn^2+^SOD (PDB ID 8VHY, solved at 2.3 Å resolution). For **e-f**, dashed lines indicate hydrogen bonds ≥ 1.8 Å, and hashed lines indicate SSHBs that are hydrogen bonds < 1.8 Å.

Trp161Phe MnSOD, like Tyr34Phe MnSOD, exhibits kinetics indicating a highly product-inhibited enzyme (*k*_3_ >> *k*_2_, **Table 1**). HERFD-XANES data of reduced and peroxide-soaked Trp161Phe MnSOD demonstrates the same trends as Tyr34Phe MnSOD, with a lower intensity at ∼6550 eV and a higher intensity at ∼6562-6570 eV (**Fig. 4b**). Likewise, a lower intensity for the pre-edge maxima at ∼6540 eV and a greater intensity for the pre-edge tail at ∼6542.5 eV is seen for peroxide-soaked Trp161Phe MnSOD compared to the reduced counterpart. Overall, similar trends are observed between reduced and peroxide-soaked forms of Tyr34Phe and Trp161Phe MnSOD.

Next, we wondered how well the product-inhibited complex could be isolated in wildtype MnSOD, where *k*_3_ = *k*_2_ (**Table 1**). Compared to Tyr34Phe and Trp161Phe, wildtype has the least noticeable difference in intensity among the 6550-6560 eV region between its peroxide-soaked and reduced counterparts (**Fig. 4c**). However, a higher intensity at ∼6570 eV is still observed for peroxide-soaked wildtype as well as the higher intensity for the pre-edge at ∼6542.5 eV. Of the reduced and peroxide-soaked pairs, the spectra of the wildtype pair are the most similar in their overall shape. Since the wildtype enzyme exhibits physiological product inhibition levels (*k*_3_ = *k*_2_, **Table 1**) compared to the Tyr34Phe and Trp161Phe variants that are enriched for product inhibition (*k*_3_ >> *k*_2_ and lower *k*_4_ compared to wildtype, **Table 1**), a potential explanation for the less pronounced differences for the wildtype pair is that the peroxide-soaked data may reflect a mixture of species rather than a fully isolated product-bound complex.

When comparing the HERFD-XANES data of wildtype, Tyr34Phe, and Trp161Phe peroxide-soaked MnSOD variants, several commonalities and trends are observed. First, the edge of the three samples lay on top of each other (**Fig. 4d**), indicating that the oxidation and spin state is maintained across the variants. For the pre-edge, the apex of the pre-edge intensity for the peroxide-soaked samples is found to be consistently lower than the reduced counterparts in addition to an increase in the intensity of the high-energy pre-edge tail (inset, **Fig. 4a-d**). While the apex peak of the pre-edge shifts within 0.2 eV among the samples tested (inset, **Fig. 4d**), the significance of the shift is unclear due to the closeness of the peaks. For the overall spectral shape, peroxide-soaked Trp161Phe MnSOD has the most drastic shift compared to its reduced counterpart (6550-6580 eV, **Fig. 4b**). Furthermore, peroxide-soaked Trp161Phe MnSOD also has the lowest intensity at the 6550 eV region compared to other peroxide-soaked variants (**Fig. 4d**). This may be because Trp161Phe MnSOD is the most product-inhibited of the three MnSOD forms tested, (*k*_3_ >> *k*_2_ and lowest *k*_4_, **Table 1**). Wildtype is the least product-inhibited when compared to Trp161Phe and Tyr34Phe, (*k*_3_ = *k*_2_ and higher *k*_4_, **Table 1**) and has the least drastic shape shift between its reduced and peroxide-soaked counterparts. Tyr34Phe, while still highly product-inhibited (*k*_3_ >> *k*_2_), has a *k*_4_ value between that of Trp161Phe and wildtype and may explain why the 6550 eV intensity is intermediate (**Fig. 4d**). While previous kinetic models ascribe *k*_4_ as the zero-order decay of the inhibited complex to a trivalent Mn ion^33–35,66,92^, some studies have indicated that the product-inhibited complex decays to divalent Mn ion^66,67^. Indeed, exposing our samples to lesser amounts of peroxide before HERFD-XANES data collection leads to spectra that more closely resemble reduced spectra (**Supplementary Fig. 2**). These observations indicate that retention of the inhibited complex is not exclusively dependent on a proton transfer from Tyr34 but instead on other factors of the active site.

To provide insight into the mechanism of product inhibition, we compared our D_2_O_2_-soaked Tyr34Phe MnSOD neutron structure with that of our previous D_2_O_2_-soaked Trp161Phe MnSOD neutron structure (**Fig. 4e**, PDB ID 8VHW)^70^. The orientation of the singly-protonated dioxygen ligand differs between the two structures. The O^1^-O^2^(LIG) axis for Trp161Phe is nearly coaxial with that of the Mn-Asp159 bond. For Tyr34Phe, the O^1^-O^2^(LIG) axis is at an angle of ∼55° to the Mn-Asp159 bond. These orientation differences are most likely attributed to the residue at position 161. LIG closely interacts with Trp161 in Tyr34Phe MnSOD (**Fig. 3a**), and when this contact is absent, LIG can assume a different orientation. Interestingly, the strong LIG- Gln143 hydrogen bond in Trp161Phe MnSOD coincides with a longer retention of the product inhibited- complex (*k*_4_, **Table 1**). However, since the 161 position is occupied by a tryptophan residue in wildtype, the LIG orientation in Tyr34Phe MnSOD is probably close to what would occur physiologically.

Both Tyr34Phe and Trp161Phe MnSOD have nearly ablated catalysis for the Mn^2+^ to Mn^3+^ redox reaction (*k*_2_, **Table 1**) and therefore proceed predominately through the product-inhibited pathway (*k*_3_ and *k*_4_, **Table1**). To investigate whether deficient *k*_2_ catalysis in Tyr34Phe is from a loss of the Tyr34 ionizable group or the distortion of the Gln143 position, we compared the neutron structure of Tyr34Phe Mn^2+^SOD with our previous Trp161Phe Mn^2+^SOD structure that preserves the number of ionizable groups (**Fig. 4f**, PDB ID 8VHY)^70^. Both variant structures have a lengthened WAT1-Gln143 interaction compared to wildtype which is a site of proton transfer (**Fig. 1b**)^13^. This suggests that the *k*_2_ PCET reaction is not solely dependent on the ionization of Tyr34 but also on tight WAT1-Gln143 hydrogen bonding. Furthermore, the weakened interaction for the variants allows WAT1 to be more easily displaced by dioxygen species for the formation of the product-inhibited complex through *k*_3_ (**Table 1**). Altogether, both the Tyr34Phe and Trp161Phe Mn^2+^SOD structures highlight the importance of the WAT1-Gln143 interaction for catalysis.

Previous studies have indicated that relief of inhibited complex (i.e., disassociation of the dioxygen species from the Mn ion) is proton transfer dependent, with Tyr34 being the most obvious proton donor^33,34^. Indeed, mutation of Tyr34 to phenylalanine leads to a slower disassociation of the inhibited complex (*k*_4_, **Table 1**). However, several other point mutations lead to the same effect, including Trp161Phe and Trp123Phe^33,34^. What these variants have in common with Tyr34Phe is that the affected residue position is directly adjacent to Gln143, which has been shown to change protonation states (**Fig. 1b**)^13^. From the D_2_O_2_-soaked neutron structure of Trp161Phe MnSOD, we previously postulated that the strong interaction between LIG and Gln143 could represent a possible proton transfer site (**Fig. 4e**)^70^. However, this suggestion is at odds with the Tyr34Phe counterpart, which has a weak hydrogen bonding interaction with the already protonated O^2^(LIG) atom and a faster Mn-dioxo dissociation (*k*_4_, **Table 1**). This means Gln143 may not directly protonate LIG and that a stronger LIG-Gln143 interaction instead contributes to longer retention of the inhibited complex. An alternative proton donor for the protonation of LIG to form H_2_O_2_ could be a solvent molecule. Proton donation from a solvent molecule would have to compete with the LIG-Gln143 interaction, and this also explains why a short LIG-Gln143 hydrogen correlates with longer retention of the inhibited complex. Overall, our analysis of D_2_O_2_-soaked MnSOD variants suggests that Gln143 plays a role in the retention of the inhibited complex.

### The electronic configuration of the Mn ion

For enzymes that use metal centers to catalyze redox reactions, the arrangement of the metal 3d orbitals determines how electrons are exchanged and how substrates orient for catalysis^93^. For MnSOD, the metal ion is in a distorted *C*_3*v*_ symmetry environment with either 4 or 5 occupied electrons in the α-manifold, depending on the oxidation state of the metal^76^. For Mn^3+^ with S = 2, the e_π_ (xz/yz) and e_σ_ (xy/x^2^-y^2^) α orbitals are occupied, while Mn^2+^ with S = 5/2 also has the z^2^ α orbital occupied (**Fig. 5a**). The z^2^ α orbital exchanges electrons during redox reactions as it is the lowest unoccupied molecular orbital (LUMO) for Mn^3+^ and the highest occupied molecular orbital (HOMO) for Mn^2+^. This means that mono- or dioxygen species are most likely to bind the Mn ion along the z-axis for reactivity (**Fig. 5b**). For further insight into the MnSOD metal 3d orbitals, we compared the K-pre-edge spectra found through time-dependent DFT (TD-DFT) simulations with spectra measured from HERFD-XANES.

**Fig. 5:**
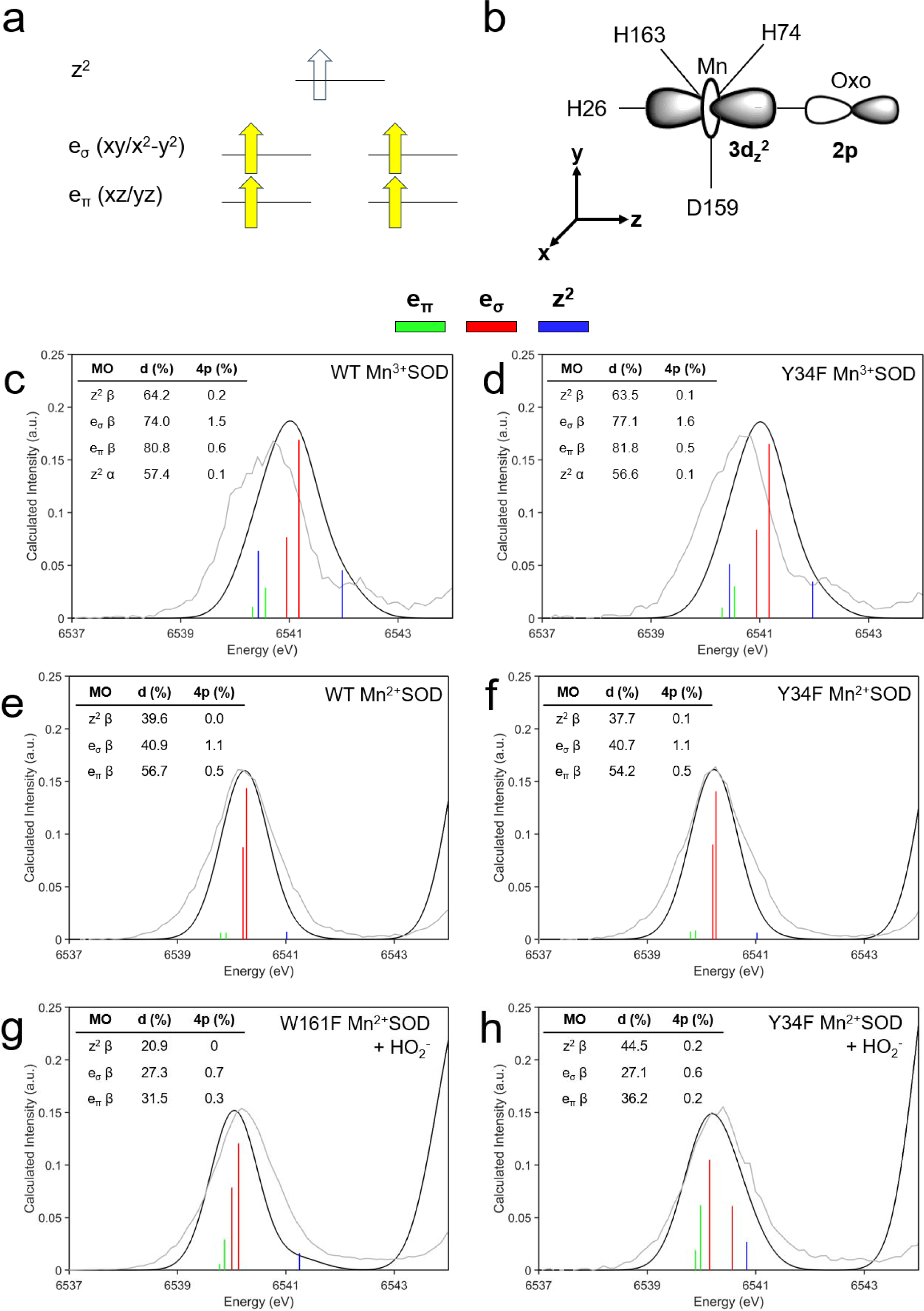
MnSOD metal 3d orbital transitions. **a** Schematic molecular orbital diagram of high spin Mn^3+^ and Mn^2+^ complexes. The e_σ_ and e_π_ orbitals of the α-manifold are occupied for Mn^3+^, while Mn^2+^ also has the z^2^ α orbital occupied. The z^2^ α orbital exchanges electrons during catalysis. **b** Schematic of the Mn ion in an idealized *C*_3*v*_ environment highlighting the interaction of the Mn ion 3d_z_^2^ orbital with the 2p orbital of a mono- or dioxygen species. **c-h** Comparison of experimental MnSOD K-pre-edge spectra with TD-DFT simulated intensities. The metal 3d orbital transitions are colored by vertical lines: e_π_ in green, e_σ_ in red, and z^2^ in blue. The tables within the panels list the percentage 3d and 4p molecular orbital character found from ground-state DFT and Löwdin analysis. Bond distances of DFT-optimized structures are found in **Supplementary Table 4**.

In K-edge XAS, the pre-edge corresponds to 1s to 3d transitions that gain intensity through 4p character mixing into the 3d orbitals from symmetry distortions and/or loss of inversion symmetry^73–75^. The contribution of the 4p electric dipole is about one hundred times stronger than the 3d quadrupole contributions, so a small amount of 4p mixing can significantly influence the pre-edge spectra^73,74,77^. With TD-DFT simulations, the 3d/4p contributions can be assigned to the experimental pre-edge spectra (**Fig. 5c-h**). In the plots, the vertical stick heights represent the intensity of 1s to 3d/4p transitions from quadrupole and dipole contributions, and percentage of 3d and 4p character from ground-state DFT are listed. To identify the impact of the Tyr34Phe variant on the electronic configuration of the metal, we compared its Mn^3+^SOD, Mn^2+^SOD, and H_2_O_2_-soaked pre-edge spectra to those of wildtype and Trp161Phe.

The experimental pre-edge spectra are similar between wildtype and Tyr34Phe Mn^3+^SOD, indicating that the Tyr34Phe variant does not significantly alter the Mn^3+^ ion orbital configuration (**Fig. 5c, d**). The TD-DFT spectra are also similar between the two complexes, though the simulated spectra underestimate the splitting of orbital transitions and overestimates intensity. This misestimation has been previously reported for other transition metal complexes of similar symmetry^77,94^. In brief, TD-DFT overestimates the exchange interaction between the 1s core hole and the valence 3d orbitals, which results in decreased energy splitting between transitions. Regardless, the e_σ_ orbitals are the primary contributors to the intensities of the pre-edge as they have significant dipole character from 4p mixing. Indeed, ground-state DFT indicates that the e_σ_ β orbitals have 1.5-1.6% 4p character and have the highest 4p mixing compared to the other 3d orbitals (inset table, **Fig. 5c, d**). In the experimental spectra, a high-energy tail is observed at ∼6542.5 eV, which is likely to be the z^2^ β transition as indicated by TD-DFT. Assignment of the lower energy z^2^ α and e_π_ β transitions in the experimental spectra is less clear, but they may contribute to the left shoulders of the pre-edge peaks. Despite the misestimation of the energy splitting with TD-DFT, the contributions of the e_σ_ and z^2^ β transitions can be assigned to the experimental spectra.

For wildtype and Tyr34Phe Mn^2+^SOD, the experimental pre-edge spectra are nearly identical (**Fig. 5e, f**). DFT calculations suggest that the e_σ_ β orbitals contribute the majority of the intensity to the pre-edge for both wildtype and Tyr34Phe Mn^2+^SOD due to 4p mixing. The z^2^ β transitions found at higher energies are of pure quadrupole character that contributes weak intensity. At lower energy, the e_π_ β transitions likely contribute weak intensity to the left shoulders of the pre-edge peaks. For the resting state Mn^2+^SOD complexes, the experimental and simulated spectra agree with each other and allow the assignment of all the orbital transitions.

The H_2_O_2_-soaked Trp161Phe and Tyr34Phe complexes have similar experimental spectra though the calculated spectra suggest differences in orbital transitions (**Fig. 5g, h**). The calculated spectra used a Mn^2+^ ion along with a HO_2_^−^ as the dioxygen species. For Trp161Phe, the e_σ_ and e_π_ β transitions are close in energy and suggest further distortions of *C*_3*v*_ symmetry and mixing of the d orbitals (**Fig. 5g**). These e transitions are mostly of dipole character, with the e_σ_ set bearing the most. The z^2^ β orbital is the highest energy transition with weak intensity. For Tyr34Phe, the d orbitals also mix, and the e_σ_ transitions begin to split (**Fig. 5h**). Between Tyr34Phe and Trp161Phe, the energy of the z^2^ β transition differs, with that of Trp161Phe being ∼0.4 eV higher than Tyr34Phe (**Fig. 5g, h**). This is reflective of different Mn ion interactions along the z-axis, like with HO_2_^-^. However, the calculated intensities of the z^2^ β transitions are weak, and the energies that these transitions occur in the experimental spectra are not obvious. Overall, the pre-edge spectral shapes of H_2_O_2_-soaked Trp161Phe and Tyr34Phe complexes are similar, though they have different transition energies.

Analysis of the K-pre-edge for various MnSOD complexes indicates that the hydroxyl group of Tyr34 does not significantly affect the metal 3d transitions of the Mn^3+^SOD and Mn^2+^SOD resting states (**Fig. 5c-f**). The pre-edge spectra for wildtype and Tyr34Phe resting states are dominated by e_σ_ transitions that are flanked by weaker intensity e_π_ and z^2^ transitions. The majority of the intensities from the e_σ_ orbitals are from dipole contributions as a result of 4p mixing. Specifically, the metal 4p_x_ and 4p_y_ mix with the 3d_x_^2^-_y_^2^ and 3d_xy_ orbitals. These orbitals have σ-overlap with the Mn ligands along the xy plane, namely with residues His74, His163, and Asp159 (**Fig. 5b)**. The deviation of these trigonal ligands away from idealized 120° angles in *C*_3*v*_ symmetry results in 4p mixing and significant intensity contributions to the pre-edge peak^77^.

The TD-DFT calculations for the Trp161Phe and Tyr34Phe Mn^2+^SOD that are bound by HO_2_^-^ have similar spectral shapes though different energies of for the e_σ_ and z^2^ transitions (**Fig. 5g, h**). This may, in part, be explained by the different orientations of HO_2_^−^ bound to the Mn^2+^ ion that leads to different orbital characteristics (**Fig. 4e**). The orbital transitions for these complexes have strong covalent mixing with the Mn-bound ligands, leading to less overall d character compared to the resting state complexes. The d orbitals also mix with each other and suggest the complexes are further distorting away from *C*_3*v*_ symmetry (**Fig. 5g, h**). TD-DFT analysis of dioxygen-bound Trp161Phe and Tyr34Phe Mn^2+^SOD suggest that different modes of HO_2_^-^ binding may result in similar spectra.

### Tyr34 contributes to the pK_a_s of Tyr166/His30 and influences the proton shuffle across the subunit interface

Second-sphere residues His30 and Tyr166 have unusual pK_a_s in wildtype MnSOD and change protonation states (**Fig. 1c**)^13^. To identify whether the Tyr34 mutation to Phe affects the protonations of His30/Tyr166, we investigated the nuclear density maps of Tyr34Phe MnSOD in D_2_O_2_-soaked, reduced, and oxidized forms. For chain A of D_2_O_2_-soaked MnSOD that has dioxygen species bound (**Fig. 3a, b**), the omit |*F*_o_| - |*F*_c_| density indicates that Tyr166 is in the neutral, protonated form while His30 is singly protonated on the N^δ1^ atom (**Fig. 6a**). Interestingly, for chain B, Tyr166 is instead deprotonated and His30 is now protonated on N^ε2^ (**Fig. 6b**). The two residues form a 1.7 Å SSHB due to the negative charge of ionized Tyr166. Different chains of the same MnSOD neutron structure have been observed before to differ in protonation states, with the most explicable cause being differences in solvent accessibility^13^. However, the protonation configuration observed in chain B is unique and has not been seen before (**Fig. 6b**). It is not known if this proton configuration occurs in wildtype enzyme as neutron data for D_2_O_2_-soaked wildtype has yet to be collected. But we can compare reduced and oxidized Tyr34Phe MnSOD to wildtype^13^.

**Fig. 6:**
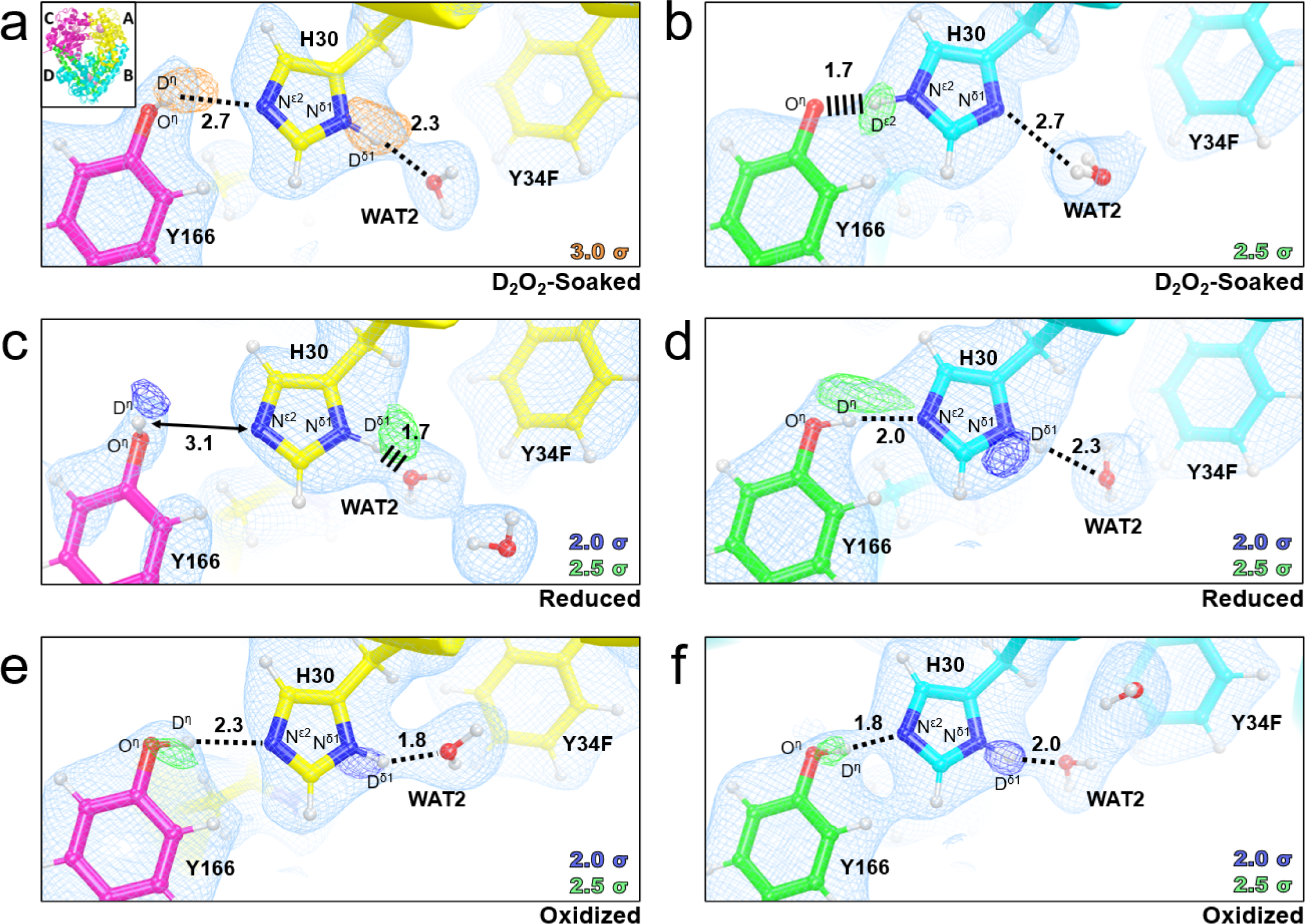
Protonation states of Tyr166 and His30 in Tyr34Phe MnSOD. **a-b** Neutron structures of D_2_O_2_-soaked Tyr34Phe MnSOD (2.3 Å resolution). **c-d** Neutron structures of Tyr34Phe Mn^2+^SOD (2.5 Å resolution). **e-f** Neutron structures of Tyr34Phe Mn^3+^SOD (2.3 resolution). Blue, green, and orange omit |*F*_o_| - |*F*_c_| difference neutron scattering length density of protons displayed at 2.0 σ, 2.5σ, and 3.0σ, respectively. Light blue 2|*F*_o_| - |*F*_c_| density is displayed at 1.0σ. Distances are in Å. Dashed lines indicate hydrogen bonds ≥ 1.8 Å and hashed lines indicate SSHBs that are hydrogen bonds < 1.8 Å. Chains are colored according to the inset in Fig. 6a.

For reduced Tyr34Phe MnSOD, both chains have the same protonation states (**Fig. 6c, d**). Tyr166 is protonated and neutral while His30 is singly-protonated on the N^δ1^ atom. The omit density for D^η^(Tyr166) is elongated and spans to N^ε2^(His30) (**Fig. 6d**). Elongated density has been observed between the two residues before in reduced wildtype MnSOD^13^. This may be indicative of a proton transfer occurring between Tyr166 and His30, in line with the different protonation states observed in the D_2_O_2_-soaked structure (**Fig. 6a, b**).

The protonation states of oxidized Tyr34Phe (**Fig. 6e, f**) are identical to that of reduced Tyr34Phe (**Fig. 6c, d**), which is in contrast to wildtype MnSOD where the His30 protonation alters between oxidized and reduced states (**Fig. 1c**). This difference is probably due to the Phe mutation at position 34. In wildtype, Tyr34 is ionized when the Mn ion is oxidized (**Fig. 1d**). The negative charge on Tyr34 may exert electronic effects that alter the pK_a_ of His30. Thus, Tyr34 may help modulate the pK_a_ of nearby residues so efficient proton transfers occur for PCET catalysis.

Our three neutron structures of Tyr34Phe MnSOD highlight the importance of residues Tyr166, His30, and Tyr34 in maintaining the proton pool of the active site. Two protons are needed to protonate O_2_^2−^ to H_2_O_2_ during the fast Mn^2+^ → Mn^3+^ reaction (*k*_2_, **Table 1**). The D_2_O_2_-soaked structure unambiguously indicates that His30 and Tyr166 shuffle protons, with proton transfers between O^η^(Tyr166) and N^ε2^(His30) coinciding with changes in the N^δ1^(His30) protonation state (**Fig. 6a, d**). In wildtype MnSOD, the proton between O^η^(Tyr166) and N^ε2^(His30) was instead observed to be shared, perhaps because the wildtype structures were collected at room temperature (**Fig. 1c, d**). Another distinction between Tyr34Phe and wildtype MnSOD is that in the oxidized forms, the N^δ1^(His30) have different protonation states. For Tyr34Phe, N^δ1^(His30) is protonated (**Fig. 6e, f**) while in wildtype N^δ1^(His30) is deprotonated (**Fig. 1c**). The pK_a_s of active site residues are a result of multiple effects, including residue composition and ionization states. An ionized Tyr34 coincides with a deprotonated N^δ1^(His30) in wildtype MnSOD and suggests that Tyr34 contributes to modulating the pK_a_ of nearby residues for oxidized MnSOD. Overall, the Tyr34Phe MnSOD neutron structures help elucidate the role of Tyr166, His30, and Tyr34 in MnSOD catalysis.

### Summary and comparison of Tyr34Phe active site configurations

Comparing the Tyr34Phe MnSOD neutron structures with our previous wildtype and Trp161Phe neutron structures helps our understanding of the role of Tyr34 in catalysis. For oxidized wildtype and Tyr34Phe MnSOD, the WAT1-Gln143 protonation states and hydrogen bond interaction are similar (**Fig. 7a, b**). However, in reduced Tyr34Phe, His30 is observed to have different protonation states compared to wildtype, and the proton between His30 and Tyr166 is not shared. For the reduced counterparts, the hydrogen bonding of Gln143 with WAT1 and Trp123 is weaker in Tyr34Phe MnSOD (**Fig. 7c, d**). The Gln143 hydrogen bonds in wildtype are SSHBs (< 1.8 Å), while those of Tyr34Phe are typical hydrogen bonds (> 1.8 Å). Reduced wildtype is observed to have a shared proton between His30 and Tyr166 while Tyr34Phe does not. However, chain B of reduced Tyr34Phe MnSOD is observed to have elongated nuclear density between His30 and Tyr166 (**Fig. 6d**), which indicates that a proton may be shared. For peroxide-bound Trp161Phe and Tyr34Phe, HO_2_^-^ binds the Mn ion in different orientations (**Fig. 7e, f**). Furthermore, in Trp161Phe, a H_2_O_2_ molecule is bound with SSHBs between His30 and an anionic Tyr34. The lack of H_2_O_2_ at this site in Tyr34Phe may be due to the absence of the hydroxyl group. Overall, Tyr34 plays a role in several aspects of the active site, and several enzymatic details are revealed from this collection of neutron structures.

**Fig. 7:**
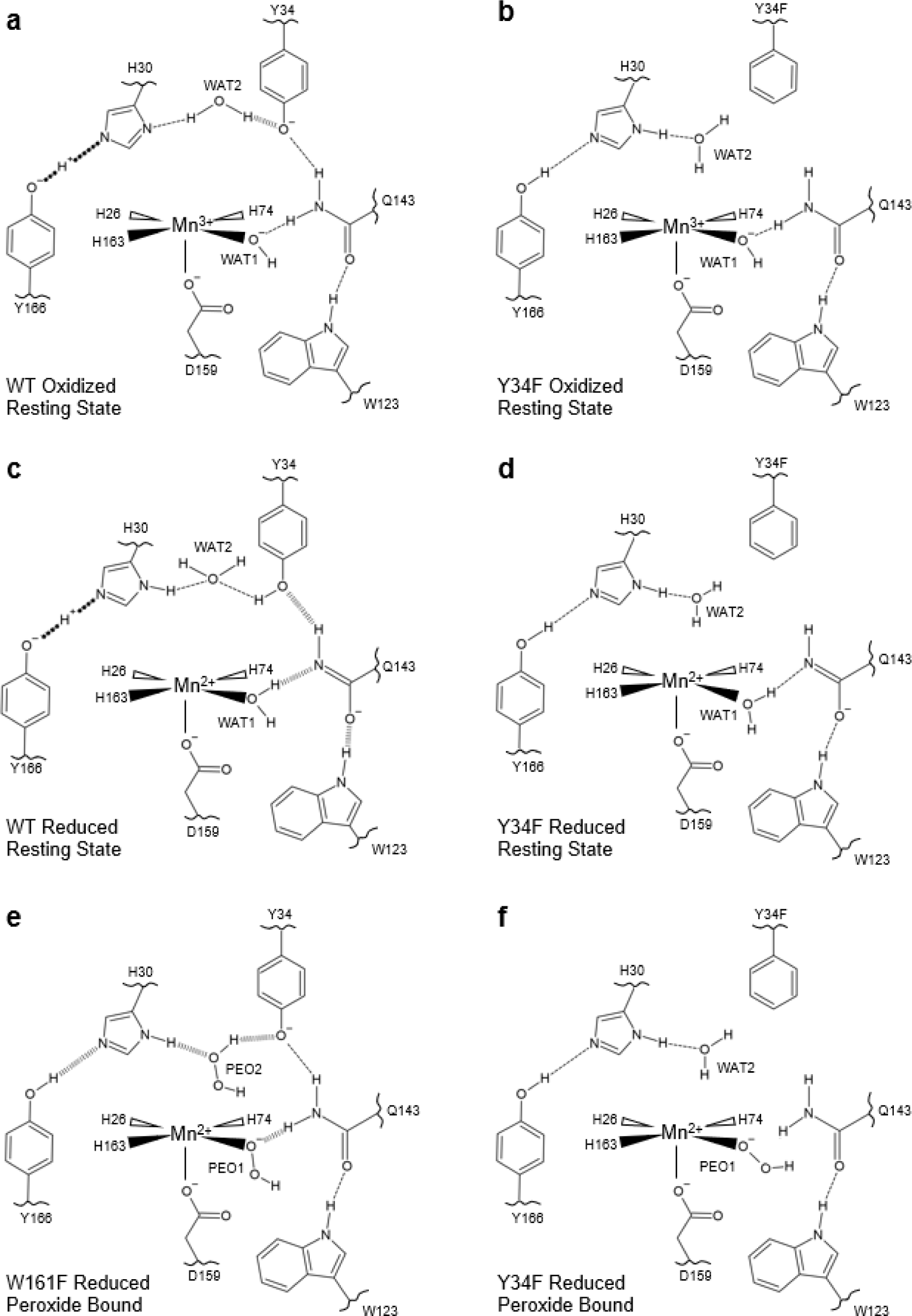
Comparison of Tyr34Phe MnSOD active site configurations to wildtype and Trp161Phe. **a** Wildtype oxidized resting state from chain B of PDB ID 7KKS^13^. **b** Tyr34Phe oxidized resting state from chain A (present study, Fig. 3e**, 6e**). **c** Wildtype reduced resting state from chain B of PDB ID 7KKW^13^. **d** Tyr34Phe reduced resting state from chain A (present study, Fig. 3c**, 6c**). **e** The product-inhibited complex of Trp161Phe, consisting of a HO2 molecule bound to a reduced active site. From chain B of PDB ID 8VHW^70^. **f** The product-inhibited complex of Tyr34Phe, consisting of a HO_2_^-^ molecule bound to a reduced active site. From chain A of the present study (**Fig. 3a-b, 6a**). Dashed lines represent normal hydrogen bonds (> 1.8 Å), wide-dashed represent SSHBs (hydrogen bonds <1.8 Å), and round-dotted lines represent a shared proton. The portrayal of hydrogen bond lengths in 2D are not representative of those seen experimentally in 3D.

First, Tyr34 helps control the charge at the active site and provides an environment conducive to charge-dependent proton and electron transfers. For example, Tyr34 and His30 both lose protons when the active site is oxidized and gain protons when the active site is reduced (**Fig. 7a, 7c, 1c-d**). As a result, we previously hypothesized that Tyr34 and His30 protonate O_2_^2-^ during the fast Mn^2+^ → Mn^3+^ reaction (*k*_2_, **Table 1**)^13^. In the Tyr34Phe variant, the His30 protonation state remains consistent between resting states, which implicates Tyr34 in influencing the pK_a_ of His30 through electronic effects (**Fig. 7b, 7d, 6c-f**). These charge effects of Tyr34 may be a partial contributor to the deficient catalysis observed for the Tyr34Phe variant (**Table 1**).

Second, Tyr34 plays a significant role in the fast Mn^2+^ → Mn^3+^ reaction as suggested by the nearly ablated *k*_2_ for Tyr34Phe (**Table 1**). Comparison of the resting Mn^2+^SOD neutron structures of wildtype and Tyr34Phe indicates that Tyr34 enforces a strong WAT1-Gln143 interaction (**Fig. 7c-d, 3d**). This interaction is important for PCET, where the WAT1 → Gln143 proton transfer coincides with the Mn^2+^ → Mn^3+^ redox reaction (**Fig. 1b**)^13^. Another potential reason for ablated *k*_2_ catalysis for the Tyr34Phe variant is the loss of an ionizable group that could protonate O_2_^2-^ or HO_2_^-^. However, the Trp161Phe variant also has an ablated *k*_2_ and maintains the number of ionizable groups (**Table 1**). The shared feature for these two variants is the weakened WAT1-Gln143 interaction (**Fig. 4f**). Since our previous structures indicate that Tyr34 loses a proton during the Mn^2+^ → Mn^3+^ reaction (**Fig. 1d**), it is possible that the WAT1 → Gln143 proton transfer precludes a Tyr34 → O_2_^2−^ proton transfer. Overall, our structures indicate Tyr34 contributes to orienting WAT1 and Gln143 for a proton transfer event during the Mn^2+^ → Mn^3+^ redox reaction.

Third, the formation of the product-inhibited complex is dependent on Tyr34. The complex is characterized by an HO_2_^−^ molecule replacing the WAT1 position in the Mn^2+^ oxidation state (**Fig. 7e-f, 4e**). As such, the capacity to form the inhibited complex may correlate with the ease an HO_2_^-^ molecule can displace WAT1. The Tyr34Phe and Trp161Phe variants have a higher propensity to accumulate the inhibited complex, and both have a weakened WAT1-Gln143 interaction in the Mn^2+^ oxidation state that would permit easier displacement of WAT1 (**Fig. 4f**). This suggests that the Tyr34-Gln143-WAT1 hydrogen bond network suppresses product inhibition.

Lastly, retention of the product-inhibited complex correlates with the strength of hydrogen bonding between HO_2_^−^ and Gln143. The inhibited complexes captured in the Tyr34Phe and Trp161Phe variants reveal two different HO_2_^−^ binding orientations and hydrogen bond interactions (**Fig. 7e-f, 4e**). The hydrogen bonding between HO_2_^−^ and Gln143 is stronger in Trp161Phe compared to Tyr34Phe, and this stronger interaction correlates with a slower disassociation of the inhibited complex for Trp161Phe (*k*_4_, **Table 1**). As protonation of HO_2_^−^ to H_2_O_2_ is required for relief of the inhibited complex, the hydrogen bonding of Gln143 with HO_2_^−^ may compete with a H_2_O → HO_2_^−^ proton transfer to form H_2_O_2_. Comparison of the product-inhibited complex of Tyr34Phe and Trp161Phe provides clues into the relief of the inhibited complex.

### Suggested mechanism

From our collection of neutron structures and XAS data, we have constructed a mechanism for the fast reaction pathways *k*_1_ and *k*_2_ and the reversible product inhibition reaction pathways *k*_3_ and *k*_4_ (**Fig. 8**). Note that our data describe the inhibited complex as Mn^2+^-containing (**Fig. 2a, 4a-c**), in contrast with several mechanistic models that assign a Mn^3+^ oxidation state to the complex^33–35,66,67^. Furthermore, in the absence of data that describe how O_2_^•−^ interacts with an oxidized active site, O_2_^•−^ reacting with the Mn^3+^ ion is represented only by Mn^3+^ gaining an electron (**Fig. 8**). For a reduced active site, O_2_^•−^ likely binds between His30 and Tyr34 for PCET to form H_2_O_2_^13,43^. Overall, the mechanism delineates how MnSOD’s reversible inhibition pathway may branch from the fast reaction pathway.

**Fig. 8:**
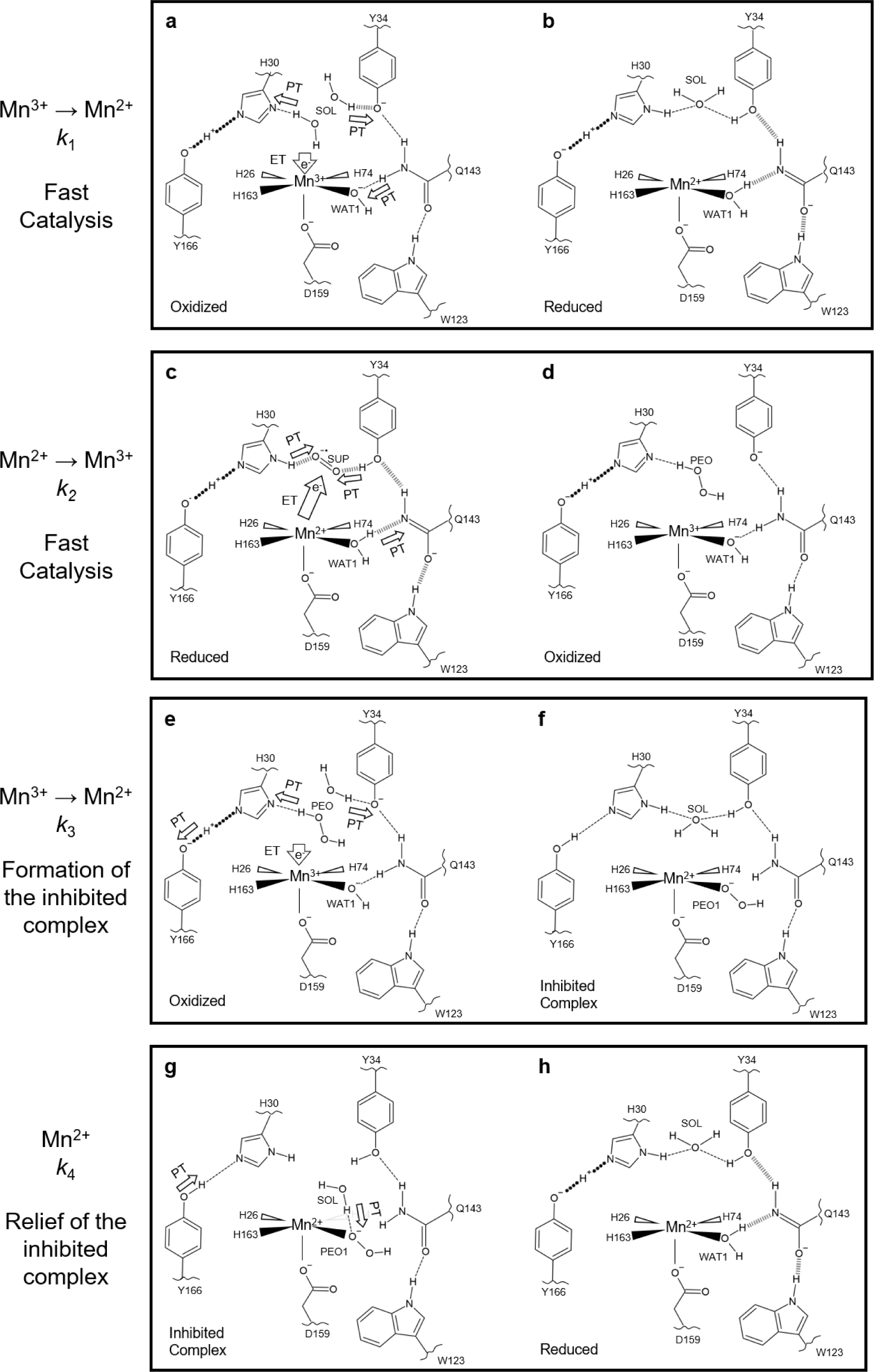
Suggested reaction mechanism of MnSOD. **a-b** The fast Mn^3+^ → Mn^2+^ half-reaction corresponding to *k*_1_. **c-d** The fast Mn^2+^ → Mn^3+^ half-reaction corresponding to *k*_2_. **e-f** Formation of the inhibited complex that is dependent on the presence of H_2_O_2_ during the reduction of the Mn^3+^ ion. In the absence of H_2_O_2_, the enzyme proceeds through the fast Mn^3+^ → Mn^2+^ half-reaction (*k*_1_). **g-h** Relief of the inhibited complex. Our data describe the inhibited complex as Mn^2+^-containing in contrast to other published mechanistic models^33–35,66,67^. In the absence of data that indicate how O_2_^•−^ interacts with an oxidized active site, O_2_^•−^ is represented only by Mn^3+^ gaining an electron. For a reduced active site, O_2_^•−^ likely binds between His30 and Tyr34 for PCET to form H_2_O_2_^13,43^. Dashed lines represent normal hydrogen bonds (> 1.8 Å), wide-dashed represent SSHBs (hydrogen bonds <1.8 Å), and round-dotted lines represent a shared proton. The portrayal of hydrogen bond lengths in 2D are not representative of those seen experimentally in 3D.

For the *k*_1_ reaction that represents the fast Mn^3+^→ Mn^2+^ half-reaction, reduction of the Mn ion occurs alongside three proton transfers, Gln143 → WAT1 and protonation of His30 and Tyr34 by solvent (**Fig. 8a**). In the reduced state, anionic Gln143 is stabilized by SSHBs with Trp123, WAT1, and Tyr34 (**Fig. 8b**). For the *k*_2_ reaction that represents the fast Mn^2+^ → Mn^3+^ half-reaction, a WAT1 → Gln143 proton transfer occurs alongside protonation and reduction of O_2_^•−^ (**Fig. 8c**). Here, O_2_^•−^ participates in a long-range electron transfer and is protonated by His30 and Tyr34 to form H_2_O_2_ (**Fig. 8d**). If Mn^3+^ gains an electron while the resultant H_2_O_2_ is still present, H_2_O_2_ donates a proton to His30 and subsequently replaces WAT1 to form the product inhibited complex (*k*_3_, **Fig. 8e-f**). The inhibited complex is characterized by the inability to perform the WAT1 → Gln143 proton transfer and prohibits fast catalysis (**Fig. 8f**). Relief of the inhibited complex is achieved by protonation of HO_2_^−^ by a solvent molecule (*k*_4_, **Fig. 8g**). H_2_O_2_ departs the active site and is replaced by WAT1 to form the Mn^2+^ resting state (**Fig. 8h**). From relief of inhibition, fast catalysis may proceed again through *k*_2_ (**Fig. 8c-d**).

The key determinant of whether product inhibition is engaged is the His30 proton donor during Mn^3+^ reduction. If H_2_O_2_ is the proton donor, the inhibited complex is formed (*k*_3,_ **Fig. 8e**). If a solvent molecule is the proton donor, the enzyme proceeds through fast catalysis (*k*_1_, **Fig. 8a**). This explains why high concentrations of O_2_^•−^ lead to product inhibition, as O_2_^•−^ may enter the active site before H_2_O_2_ departs. Tyr34 may also serve as a potential proton acceptor for H_2_O_2_ since Tyr34 also gains a proton during reduction to Mn^2+^. However, Tyr34 is not necessary to form the inhibited complex, as indicated by our Tyr34Phe neutron structure (**Fig. 3a-b**). Our work provides insights into the PCET mechanism of MnSOD, especially in the context of product inhibition.

Altogether, our investigation reveals how a single tyrosine residue has a profound effect on PCET catalysis. Tyr34 plays a part in every MnSOD kinetic step from its roles of (1) acting as a proton donor/acceptor, (2) orienting nearby molecules Gln143 and WAT1 for proton transfer, (3) limiting the formation of the inhibited complex, and (4) shortening the lifetime of the inhibited complex. These roles of Tyr34 place the residue as a central regulator of H_2_O_2_ output in the mitochondria. H_2_O_2_ produced from MnSOD has several cellular effects, including stimulating apoptotic signaling pathways^45,46^, coordinating protein localization and activity^47^, and mitochondrial biogenesis^44^. Inactivation of Tyr34 by nitration is observed in neurological disease and further highlights the physiological role of Tyr34^58–65^. In total, the work provides insight into how a PCET enzyme facilitates catalysis and how an oxidoreductase molecularly facilitates mitochondrial function.

## METHODS

### Perdeuterated expression and purification

For deuterated protein expression of MnSOD, the pCOLADuet-1 expression vector harboring full-length cDNA of *MnSOD* was transformed into *Escherichia coli* BL21(DE3) cells. Transformed cells were grown in D_2_O minimal media within a bioreactor vessel using D_8_-glycerol as the carbon source^95^. Induction was performed with 1 mM isopropyl L-D-thiogalactopyranoside, 8 mM MnCl_2_, and fed D_8_-glycerol until an OD_600_ of 15.0. Expression was performed at 37 °C for optimal Mn metal incorporation^96^. Harvested cell pastes were stored at -80 °C until purification. For protein purification (with hydrogenated reagents), cells were resuspended in a solution of 5 mM MnCl_2_ and 5 mM 3-(*N*-morpholino)propanesulfonic acid (MOPS), pH 7.8. Clarified lysate was incubated at 55 °C to precipitate contaminant proteins that were subsequently removed by centrifugation. Next, soluble protein was diluted with an equal volume of 50 mM 2-(*N*- morpholino)ethanesulfonic acid (MES) pH 5.5, yielding a final concentration of 25 mM. Measurement of pH verified a value of 5.5 after dilution. Protein was applied onto a carboxymethyl sepharose fast flow column (GE Healthcare) and eluted with a sodium chloride gradient that contained 50 mM MES pH 6.5.

### Crystallization

Perdeuterated Tyr34Phe MnSOD crystals were grown in a microgravity environment aboard the International Space Station (ISS)^97^. Crystals growth was achieved in Granada Crystallization Boxes (GCBs, Triana) through capillary counterdiffusion using fused quartz capillary tubes (VitroCom) that had inner diameters of 2.0 mm and outer diameters of 2.4 mm^98^. 25 mg ml^-1^ protein-filled capillaries were plugged with 40 mm of 2% agarose (*w*/*w*) and inserted into GCBs filled with precipitating agent composed of 4 M potassium phosphate, pH 7.8. The pH of the phosphate buffer was achieved through 91:9 ratios of K_2_HPO_4_:KH_2_PO_4_. The GCBs were delivered to the ISS by SpX-17 as part of the *Perfect Crystals* NASA payload and returned to earth 1 month later on SpX-18. The crystals within GCBs were observed to be resilient against travel damage and were placed within carry-on baggage during further aircraft travels to the UNMC Structural Biology Core Facility and ORNL. Perdeuterated Tyr34Phe MnSOD crystals were 0.3-0.6 mm^3^ in size, and further details of microgravity crystallization have been published^97^. For X-ray diffraction, crystals were grown in 1.8 M potassium phosphate, pH 7.8 by hanging-drop vapor diffusion. Protein (23 mg ml^-1^) and reservoir solution were mixed at a 1:1 ratio to give a 4.0 µL drop. Crystals for X-ray diffraction were less than 0.1 mm^3^ in size and were fully grown after 14 d.

### Crystal Manipulations

For deuterium exchange, microgravity-grown crystals were first placed in 1 mL of hydrogenated 4 M potassium phosphate pH 7.8. Deuterium was introduced with 0.1 mL incremental additions every 2 min of 4 M deuterated potassium phosphate (K_2_DPO_4_:KD_2_PO_4_) pD 7.8 (calculated by adding 0.4 to the measured pH reading) for a total of five times and a net volume addition of 0.5 mL. After 10 min, 0.5 mL of the solution was removed leading to a 1 mL solution consisting of 33% deuterium. The process was repeated enough times to gradually increase the deuterium content to ∼100%. The 4 M deuterated potassium phosphate also served as the cryoprotectant for the cryocooling process. Further details of the process were published^99^.

For redox manipulation, the deuterated potassium phosphate solutions were supplemented with either 6.4 mM potassium permanganate (KMnO_4_) to achieve the Mn^3+^ oxidation state or 300 mM sodium dithionite (Na_2_S_2_O_4_) to achieve the Mn^2+^ state. Crystals were either sealed in capillaries or in 9-well glass plates to ensure the desired oxidation state was maintained. For the Tyr34Phe structure soaked with D_2_O_2_, redox reagents were not used. The dioxygen-bound complex was achieved by supplementing the cryoprotectant that the crystal was immersed in with D_2_O_2_ at a final concentration of 1% *w*/*v* (∼0.28 M) and soaking for 1 min before cryocooling. Flash-cooling was performed with an Oxford diffraction cryostream^100^. Further details of ligand cryotrapping were published^99^.

### Crystallographic Data Collection

Time-of-flight, wavelength-resolved neutron Laue diffraction data were collected from perdeuterated crystals using the MaNDi instrument^101,102^ at the Oak Ridge National Laboratory Spallation Neutron Source with wavelengths between 2 to 4 Å. Sample sizes ranged from 0.3 to 0.6 mm^3^ and data were collected to 2.30 Å resolution for oxidized and D_2_O_2_-soaked structures, while the reduced structure was collected to 2.5 Å resolution (**Supplementary Table 5**). Crystals were held in stationary positions during diffraction, and successive diffraction frames were collected along rotations of the Φ axis. X-ray diffraction data were collected using a Rigaku FR-E SuperBright home source or the Stanford Synchrotron Radiation Lightsource (SSRL) beamline 14-1 (**Supplementary Table 5**).

### Crystallographic Data Processing and Refinement

Neutron data were integrated using the MANTID software package^103–105^ and wavelength-normalized and scaled with LAUENORM from the Daresbury Laue Software Suite^106^. X-ray diffraction data were processed using HKL-3000^107^. Refinements of both neutron and X-ray models were completed separately with PHENIX.REFINE from the PHENIX suite^108^. The refinements were intentionally performed separately due to the known perturbations that X-rays have on the solvent structure, metal redox state, and metal coordination^36,109^. The X-ray model was first refined against its corresponding data set and subsequently used as the starting model for neutron refinement. Torsional backbone angle restraints were derived from the X-ray model and applied to neutron refinement using a geometric target function with PHENIX.REFINE^108^. Mn-ligand restraints for neutron refinement were derived from DFT calculations rather than the X-ray model to remove any influence of photoreduction. The neutron refinement was performed by modeling the D atoms of the active site last to limit phase bias. Initial rounds of refinement to fit protein structure included only non-exchangeable D atoms, which have stereochemical predictable positions. Afterward, H/D atoms were modeled onto the position of each amide proton, and occupancy was refined. In general, the asymmetric units of the neutron crystal structures had a deuterium content of ∼85% for the amide backbone, and areas with low deuterium exchange (< 50 %) coincided with the presence of hydrogen bonds forming a secondary structure. Next, exchangeable proton positions of residues outside the active site (e.g., hydroxyl group of serine/tyrosine) were manually inspected for obvious positive omit |*F*_o_| - |*F*_c_| neutron scattering length density at a contour of 2.5σ or greater and modeled as a fully occupied deuterium. If the density was not obvious, and there was no chemically sensible reason for the residue to be deprotonated (which is the case for residues outside the active site), the proton position was H/D occupancy refined. D_2_O molecules outside the active site were then modeled and adjusted according to the nuclear density. Last, D atoms of the active site were modeled manually. At the active site, a residue is considered deprotonated when (1) attempts to model and refine a proton result in negative |*F*_o_| - |*F*_c_| difference neutron scattering length density, (2) all the other protons of the residue can be placed, and (3) the heavy atom that is deprotonated acts as a hydrogen-bond acceptor. As chemically ideal covalent bond distances of D atoms were ensured during model building and refinement, small deviations from D atom positions and omit |*F*_o_| - |*F*_c_| neutron scattering length density centers were expected from the data resolution (2.3-2.5 Å).

### X-ray Absorption Spectroscopy Measurements

A solution of 3 mM MnSOD (∼70 mg ml^-1^) in 25 mM potassium phosphate pH 7.8 was treated with 8 mM potassium dichromate to isolate the Mn^3+^SOD resting state, 200 mM sodium dithionite to isolate the Mn^2+^SOD resting state, and either 20 mM superoxide or 280 mM (1% *w/v*) hydrogen peroxide to isolate the product-inhibited state. Superoxide stock solutions were generated by mixing 1.3 M potassium superoxide (KO_2_, Sigma Aldridge) in a mixture of dry DMSO (Thermo Scientific) and 0.30 M di-benzo-18-crown-6-ether (Thermo Scientific) following published protocols^66,67,79,80,110^. The concentration of superoxide in the DMSO/18-crown solution was measured by a UV-Vis spectrophotometer at a wavelength of 250 nm with a molar extinction coefficient of 2686 M^-1^ cm^-1^ ^79,111^. Superoxide solutions were stored at 4 °C under a layer of argon in glass vials sealed with rubber septum.

Mn K-edge HERFD-XANES spectra were recorded at beamline 15-2 of the Stanford Synchrotron Radiation Lightsource (SSRL), while Mn K-edge EXAFS spectra were collected at beamline 7-3. At both beamlines, data were collected at 10 K using a liquid He cryostat, and the incident energy was tuned to the first derivative of an internal Mn foil at 6539 eV. X-ray irradiation was carefully monitored so that two subsequent scans of the same spot did not have photoreduction differences, and different spots along samples were scanned. When appropriate, aluminum foil was inserted into the beam path to attenuate the incident flux. For HERFD-XANES measurements, a Johann-type hard X-ray spectrometer with six Ge(333) analyzer crystals was used with a liquid-nitrogen cooled Si(311) double crystal monochromator, and energy was calibrated to a glitch with measurement of Mn Foil. For EXAFS, measurements were recorded with a 30-element Ge solid-state detector, and a Si(220) monochromator at Φ = 90° was used.

### X-ray Absorption Spectroscopy Data Analysis

EXAFS data reduction, averaging, and refinement were carried out with the LARCH software package^112^. Refinement of the *k*^2^χ(*k*) EXAFS data used phases and amplitudes obtained from FEFF^113^. For each fit, optimization of the radial distribution around the absorbing Mn ion (*r*) and the Debye-Waller factor (σ^2^) was performed. The goodness-of-it was evaluated by χ^2^ values, reduced χ^2^ values and R-factors.

### Computational Methods

All DFT calculations were performed with the ORCA quantum chemistry package version 5.0 using the B3LYP functional, the def2-TZVP basis set for all atoms, and the CPCM solvation model^114–117^. For geometry optimizations, the full active site (i.e., residues shown in the right panel of **Fig. 1a**) was included from the neutron structure where the peptide backbone was truncated and C^α^ was fixed. All atoms for residues Trp161, Phe66, and Tyr166 were fixed with the exception of hydroxyl Tyr166 proton to mimic the packing found in the native enzyme. The Mn ion used the high-spin quintet and sextet states for trivalent and divalent systems, respectively, per experimental observations^76^. A dense integration grid and tight convergence were enforced.

For TD-DFT calculations, the Mn ion was instead assigned the core property basis set, CP(PPP)^118,119^. The geometry optimized model was used and truncated to only the Mn ion and its immediate ligands. Inclusion of all active site residues for TD-DFT did not significantly alter the simulated spectra. Computed Mn K pre-edge data were plotted using a Gaussian broadening of 1 eV, and a 32.3 eV energy correction was applied in line with previous studies^73,94^.

## DATA AVAILABILITY

Coordinates and structure factors for neutron and X-ray crystallographic data presented in this study have been deposited in the Protein Data Bank. (PDB 9BVY [https://doi.org/10.2210/pdb9bvy/pdb], PDB 9BW2 [https://doi.org/10.2210/pdb9bw2/pdb], PDB 9BWM [https://doi.org/10.2210/pdb9bwm/pdb], PDB 9BWQ [https://doi.org/10.2210/pdb9bwq/pdb], and PDB 9BWR [https://doi.org/10.2210/pdb9bwr/pdb]. All relevant data supporting the key findings of this study are available within the article and its Supplementary Information files or from the corresponding author upon reasonable request.

## ACKNOWLEDGEMENTS

This research was supported by the NIH (R01-GM145647) and NASA EPSCoR (NE-80NSSC17M0030 and NE-NNX15AM82A). The UNMC Structural Biology Core Facility was funded by the Fred and Pamela Buffett NCI Cancer Center Support Grant (P30CA036727). The research at Oak Ridge National Laboratory (ORNL) Spallation Neutron Source was sponsored by the Scientific User Facilities Division, Office of Basic Energy Sciences, US Department of Energy. The Office of Biological and Environmental Research supported research at ORNL Center for Structural Molecular Biology (CSMB) using facilities supported by the Scientific User Facilities Division, Office of Basic Energy Sciences, US Department of Energy. Use of the Stanford Synchrotron Radiation Lightsource (SSRL), SLAC National Accelerator Laboratory, is supported by the US Department of Energy (DOE), Office of Science, Office of Basic Energy Sciences under Contract DE-AC02- 76SF00515. The SSRL Structural Molecular Biology Program is supported by the DOE Office of Biological and Environmental Research, and by the National Institutes of Health, National Institute of General Medical Sciences (P30GM133894). The contents of this publication are solely the responsibility of the authors and do not necessarily represent the official views of NIGMS or NIH. Quantum chemical computations were completed using the Holland Computing Center of the University of Nebraska, which receives support from the Nebraska Research Initiative.

