## Supplemental Information for "The role of Tyr34 in proton-coupled electron transfer of human manganese superoxide dismutase"

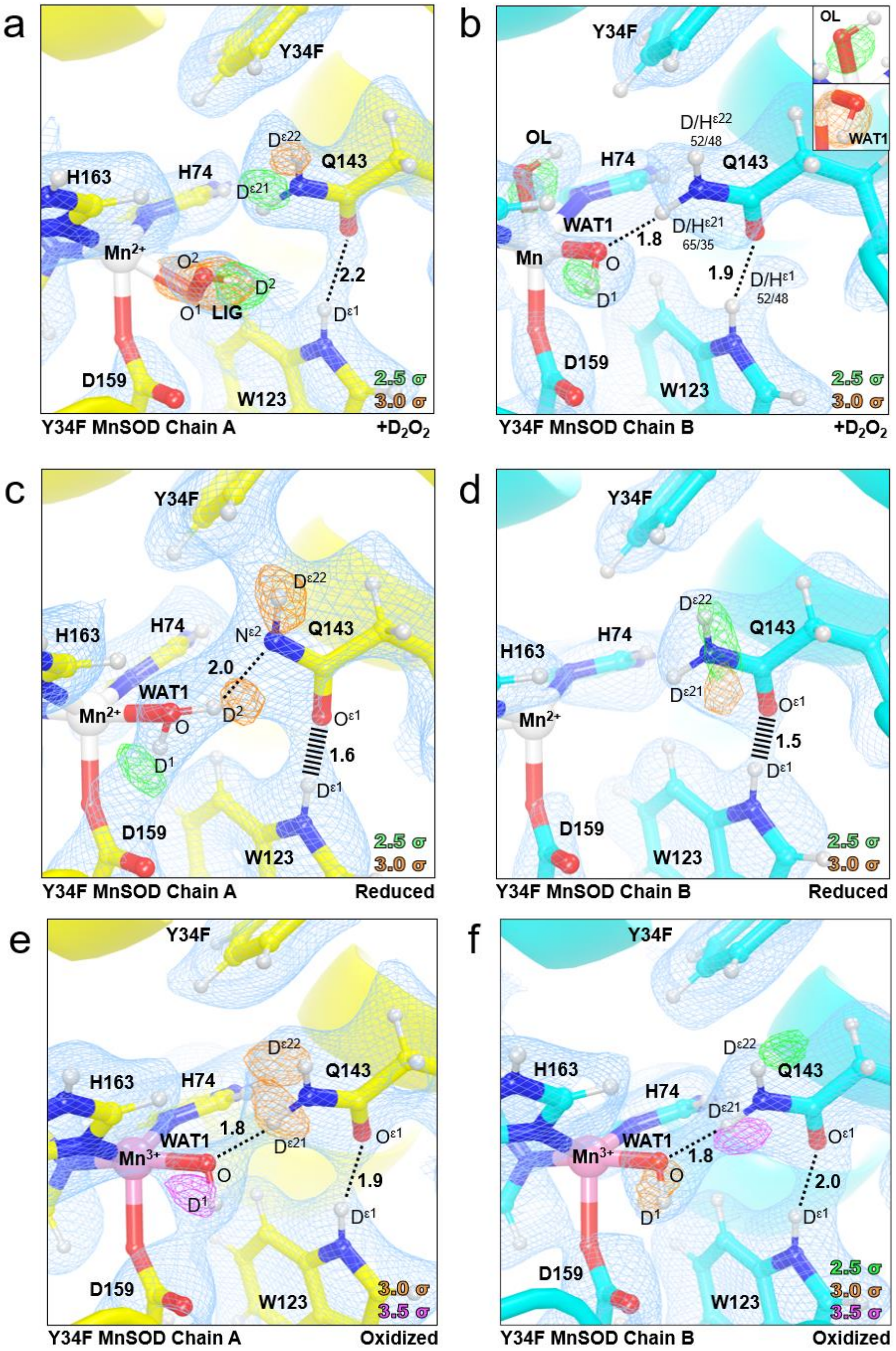


**Supplementary Figure 1: Neutron structure and protonation states at the active sites of $D_2O_2$-soaked, reduced, and oxidized Tyr34Phe MnSOD. a-b** Neutron structure of $D_2O_2$-soaked Tyr34Phe MnSOD. For the exchangeable protons of Gln143 and Trp123 of chain B, positive omit difference density was not present above 2.0σ due to density cancellation with hydrogen's negative neutron scattering length density. As an alternative, the proton positions were refined for H/D occupancy. Furthermore, an omit density for a hydroxide ion was observed to be bound to the Mn ion, opposite Asp159 (part b, inset top). WAT1 was interpreted to be a hydroxide ion (inset bottom). However, since the chain B active site contains partially-occupied atoms, all other Tyr34Phe MnSOD structures are five-coordinate, and the HERFD-XANES indicate five-coordinate complexes for Tyr34Phe MnSOD, this six-coordinate complex was not evaluated for mechanistic information. **c-d** Neutron structure of reduced Tyr34Phe MnSOD. For chain B, due to data quality issues, the nuclear density was difficult to interpret and density for WAT1 was not present. Since the HERFD-XANES data indicates that reduced Tyr34Phe MnSOD is five-coordinate like chain A, chain B was not evaluated for mechanistic information. **e-f** Neutron structure of oxidized Tyr34Phe MnSOD. Green, orange, and magenta omit $|F_o| - |F_c|$ difference neutron scattering length density of protons displayed at 2.5 σ, 3.0σ, and 3.5 σ, respectively. Light blue $2|F_o| - |F_c|$ density is displayed at 1.0σ. Distances are in Å. Dashed lines indicate typical hydrogen bonds, and hashed lines indicate SSHBs, hydrogen bonds < 1.8 Å. $D_2O_2$-soaked and oxidized structures were solved to 2.3 Å resolution, while the reduced structure was solved to 2.5 Å resolution.

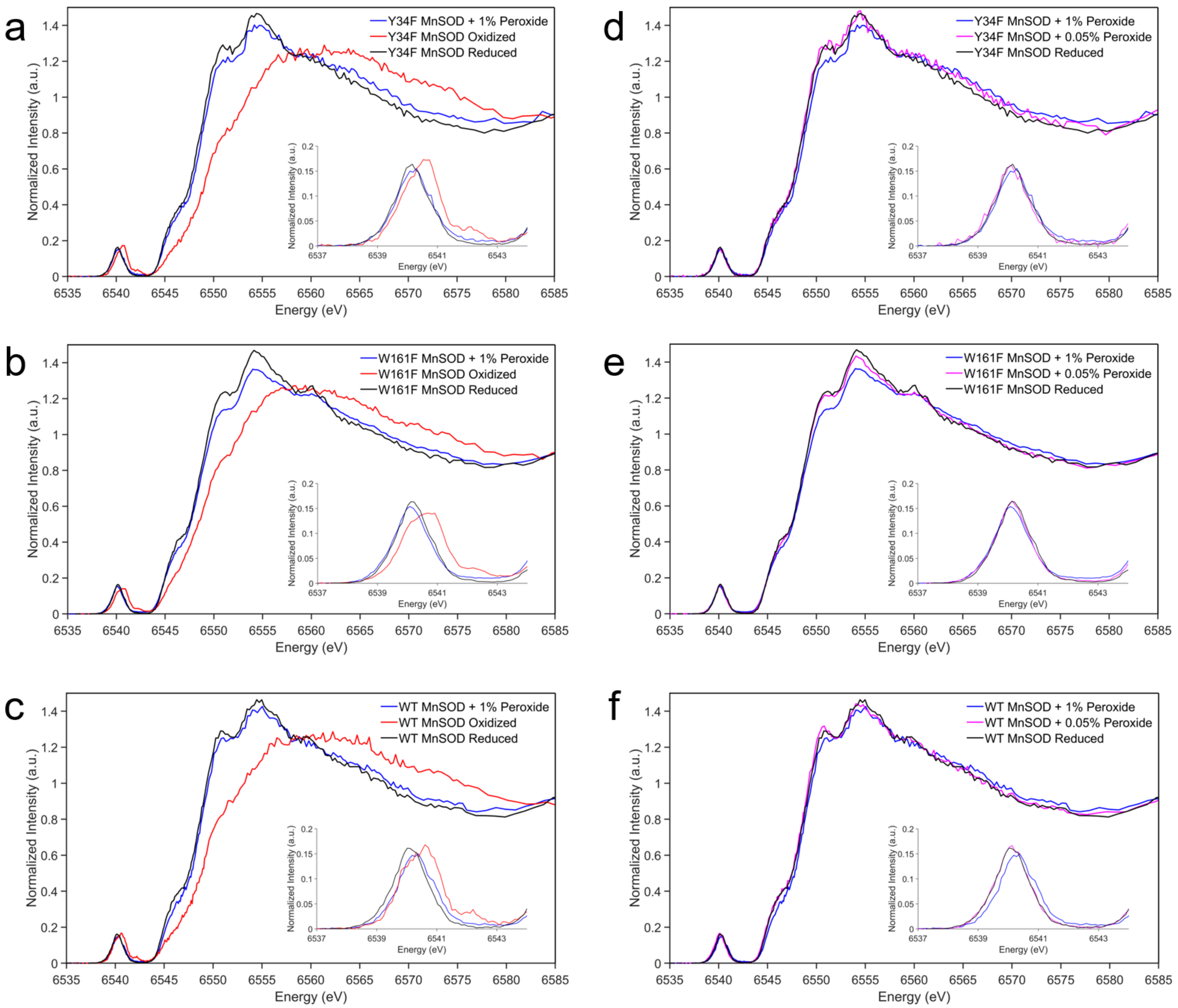


**Supplementary Figure 2: Kα HERFD-XANES spectra of MnSOD. a-c** HERFD-XANES of Tyr34Phe, Trp161Phe, and wildtype MnSOD in the 1% $H_2O_2$-soaked, oxidized, and reduced states. **d-f** HERFD-XANES of Tyr34Phe, Trp161Phe, and wildtype MnSOD in the 1% $H_2O_2$-soaked, 0.05% $H_2O_2$-soaked, and reduced states.

**Supplementary Table 1. EXAFS fitting results for superoxide-soaked Tyr34Phe MnSOD.**

**Fit with Amino Acid Ligands and Dioxygen Species (5-coordinate)**

| Mn-O | | | Mn-N | | | Mn···C | | | Mn···C | | | Mn···O | | |
|---|---|---|---|---|---|---|---|---|---|---|---|---|---|---|
| *n* | *r* (Å) | $\sigma^2$ x $10^3$ (Å$^2$) | *n* | *r* (Å) | $\sigma^2$ x $10^3$ (Å$^2$) | *n* | *r* (Å) | $\sigma^2$ x $10^3$ (Å$^2$) | *n* | *r* (Å) | $\sigma^2$ x $10^3$ (Å$^2$) | *n* | *r* (Å) | $\sigma^2$ x $10^3$ (Å$^2$) |
| 2 | 2.11 | 8.3 | 3 | 2.15 | 7.3 | 5 | 3.13 | 1.9 | 2 | 3.37 | 10 | 1 | 2.44 | 7.3 |

| Mn···O···O | | | Mn···C···O | | | Mn···C···N | | | Mn···O···C···O | | |
|---|---|---|---|---|---|---|---|---|---|---|---|
| *n* | *r* (Å) | $\sigma^2$ x $10^3$ (Å$^2$) | *n* | *r* (Å) | $\sigma^2$ x $10^3$ (Å$^2$) | *n* | *r* (Å) | $\sigma^2$ x $10^3$ (Å$^2$) | *n* | *r* (Å) | $\sigma^2$ x $10^3$ (Å$^2$) |
| 2 | 2.78 | 0.6 | 2 | 3.30 | 0 | 12 | 3.47 | 10 | 1 | 3.18 | 0 |

| $\chi^2$ | Reduced $\chi^2$ | R-Factor |
|---|---|---|
| 41.79 | 10.20 | 0.0699 |

**Fit with Amino Acids Ligands, Dioxygen Species, and Hypothetical Solvent Molecule (6-coordinate)**

| Mn-O | | | Mn-N | | | Mn···C | | | Mn···C | | | Mn···O | | |
|---|---|---|---|---|---|---|---|---|---|---|---|---|---|---|
| *n* | *r* (Å) | $\sigma^2$ x $10^3$ (Å$^2$) | *n* | *r* (Å) | $\sigma^2$ x $10^3$ (Å$^2$) | *n* | *r* (Å) | $\sigma^2$ x $10^3$ (Å$^2$) | *n* | *r* (Å) | $\sigma^2$ x $10^3$ (Å$^2$) | *n* | *r* (Å) | $\sigma^2$ x $10^3$ (Å$^2$) |
| 3 | 2.11 | 8.3 | 3 | 2.15 | 7.3 | 5 | 3.13 | 1.9 | 2 | 3.37 | 10 | 1 | 2.44 | 7.3 |

| Mn···O···O | | | Mn···C···O | | | Mn···C···N | | | Mn···O···C···O | | |
|---|---|---|---|---|---|---|---|---|---|---|---|
| *n* | *r* (Å) | $\sigma^2$ x $10^3$ (Å$^2$) | *n* | *r* (Å) | $\sigma^2$ x $10^3$ (Å$^2$) | *n* | *r* (Å) | $\sigma^2$ x $10^3$ (Å$^2$) | *n* | *r* (Å) | $\sigma^2$ x $10^3$ (Å$^2$) |
| 2 | 2.78 | 0.6 | 2 | 3.30 | 0 | 12 | 3.47 | 10 | 1 | 3.18 | 0 |

| $\chi^2$ | Reduced $\chi^2$ | R-Factor |
|---|---|---|
| 69.93 | 17.06 | 0.1170 |

**Fit Only with Amino Acid Ligands (4-coordinate)**

| Mn-O | | | Mn-N | | | Mn···C | | | Mn···C | | | Mn···O | | |
|---|---|---|---|---|---|---|---|---|---|---|---|---|---|---|
| *n* | *r* (Å) | $\sigma^2$ x $10^3$ (Å$^2$) | *n* | *r* (Å) | $\sigma^2$ x $10^3$ (Å$^2$) | *n* | *r* (Å) | $\sigma^2$ x $10^3$ (Å$^2$) | *n* | *r* (Å) | $\sigma^2$ x $10^3$ (Å$^2$) | *n* | *r* (Å) | $\sigma^2$ x $10^3$ (Å$^2$) |
| 1 | 2.11 | 8.3 | 3 | 2.15 | 7.3 | 5 | 3.13 | 1.9 | 2 | 3.37 | 10 | - | - | - |

| Mn···O···O | | | Mn···C···O | | | Mn···C···N | | | Mn···O···C···O | | |
|---|---|---|---|---|---|---|---|---|---|---|---|
| *n* | *r* (Å) | $\sigma^2$ x $10^3$ (Å$^2$) | *n* | *r* (Å) | $\sigma^2$ x $10^3$ (Å$^2$) | *n* | *r* (Å) | $\sigma^2$ x $10^3$ (Å$^2$) | *n* | *r* (Å) | $\sigma^2$ x $10^3$ (Å$^2$) |
| - | - | - | 2 | 3.30 | 0 | 12 | 3.47 | 10 | 1 | 3.18 | 0 |

| $\chi^2$ | Reduced $\chi^2$ | R-Factor |
|---|---|---|
| 54.38 | 7.661 | 0.0910 |

**Supplementary Table 2. EXAFS fitting results for peroxide-soaked Tyr34Phe MnSOD.**

**Fit with Amino Acid Ligands and Dioxygen Species (5-coordinate)**

| Mn-O | | | Mn-N | | | Mn···C | | | Mn···C | | | Mn···O | | |
|---|---|---|---|---|---|---|---|---|---|---|---|---|---|---|
| *n* | *r* (Å) | $\sigma^2$ x $10^3$ (Å$^2$) | *n* | *r* (Å) | $\sigma^2$ x $10^3$ (Å$^2$) | *n* | *r* (Å) | $\sigma^2$ x $10^3$ (Å$^2$) | *n* | *r* (Å) | $\sigma^2$ x $10^3$ (Å$^2$) | *n* | *r* (Å) | $\sigma^2$ x $10^3$ (Å$^2$) |
| 2 | 2.11 | 8.3 | 3 | 2.15 | 7.3 | 5 | 3.13 | 1.9 | 2 | 3.37 | 10 | 1 | 2.44 | 7.3 |
| Mn···O···O | | | Mn···C···O | | | Mn···C···N | | | Mn···O···C···O | | | | | |
| *n* | *r* (Å) | $\sigma^2$ x $10^3$ (Å$^2$) | *n* | *r* (Å) | $\sigma^2$ x $10^3$ (Å$^2$) | *n* | *r* (Å) | $\sigma^2$ x $10^3$ (Å$^2$) | *n* | *r* (Å) | $\sigma^2$ x $10^3$ (Å$^2$) | | | |
| 2 | 2.78 | 0.6 | 2 | 3.30 | 0 | 12 | 3.47 | 10 | 1 | 3.18 | 0 | | | |

| $\chi^2$ | Reduced $\chi^2$ | R-Factor |
|---|---|---|
| 43.08 | 10.51 | 0.0522 |

**Fit with Amino Acids Ligands, Dioxygen Species, and Hypothetical Solvent Molecule (6-coordinate)**

| Mn-O | | | Mn-N | | | Mn···C | | | Mn···C | | | Mn···O | | |
|---|---|---|---|---|---|---|---|---|---|---|---|---|---|---|
| *n* | *r* (Å) | $\sigma^2$ x $10^3$ (Å$^2$) | *n* | *r* (Å) | $\sigma^2$ x $10^3$ (Å$^2$) | *n* | *r* (Å) | $\sigma^2$ x $10^3$ (Å$^2$) | *n* | *r* (Å) | $\sigma^2$ x $10^3$ (Å$^2$) | *n* | *r* (Å) | $\sigma^2$ x $10^3$ (Å$^2$) |
| 3 | 2.11 | 8.3 | 3 | 2.15 | 7.3 | 5 | 3.13 | 1.9 | 2 | 3.37 | 10 | 1 | 2.44 | 7.3 |
| Mn···O···O | | | Mn···C···O | | | Mn···C···N | | | Mn···O···C···O | | | | | |
| *n* | *r* (Å) | $\sigma^2$ x $10^3$ (Å$^2$) | *n* | *r* (Å) | $\sigma^2$ x $10^3$ (Å$^2$) | *n* | *r* (Å) | $\sigma^2$ x $10^3$ (Å$^2$) | *n* | *r* (Å) | $\sigma^2$ x $10^3$ (Å$^2$) | | | |
| 2 | 2.78 | 0.6 | 2 | 3.30 | 0 | 12 | 3.47 | 10 | 1 | 3.18 | 0 | | | |

| $\chi^2$ | Reduced $\chi^2$ | R-Factor |
|---|---|---|
| 98.32 | 23.99 | 0.1191 |

**Fit Only with Amino Acid Ligands (4-coordinate)**

| Mn-O | | | Mn-N | | | Mn···C | | | Mn···C | | | Mn···O | | |
|---|---|---|---|---|---|---|---|---|---|---|---|---|---|---|
| *n* | *r* (Å) | $\sigma^2$ x $10^3$ (Å$^2$) | *n* | *r* (Å) | $\sigma^2$ x $10^3$ (Å$^2$) | *n* | *r* (Å) | $\sigma^2$ x $10^3$ (Å$^2$) | *n* | *r* (Å) | $\sigma^2$ x $10^3$ (Å$^2$) | *n* | *r* (Å) | $\sigma^2$ x $10^3$ (Å$^2$) |
| 1 | 2.11 | 8.3 | 3 | 2.15 | 7.3 | 5 | 3.13 | 1.9 | 2 | 3.37 | 10 | - | - | - |
| Mn···O···O | | | Mn···C···O | | | Mn···C···N | | | Mn···O···C···O | | | | | |
| *n* | *r* (Å) | $\sigma^2$ x $10^3$ (Å$^2$) | *n* | *r* (Å) | $\sigma^2$ x $10^3$ (Å$^2$) | *n* | *r* (Å) | $\sigma^2$ x $10^3$ (Å$^2$) | *n* | *r* (Å) | $\sigma^2$ x $10^3$ (Å$^2$) | | | |
| - | - | - | 2 | 3.30 | 0 | 12 | 3.47 | 10 | 1 | 3.18 | 0 | | | |

| $\chi^2$ | Reduced $\chi^2$ | R-Factor |
|---|---|---|
| 62.15 | 8.755 | 0.0753 |

**Supplementary Table 3. Active site Mn bond lengths of MnSOD neutron crystal structures.**

| Neutron | Tyr34Phe $D_2O_2$-Soaked[b] | | Tyr34Phe $Mn^{3+}$SOD | | Tyr34Phe $Mn^{2+}$SOD | | Wildtype $Mn^{3+}$SOD | | Wildtype $Mn^{2+}$SOD[c] | |
|---|---|---|---|---|---|---|---|---|---|---|
| PDB ID | 9BVY | | 9BWM | | 9BW2 | | 7KKS | | 7KKW | |
| Mn Bonds (Å) | A | B | A | B | A | B | A | B | A | B |
| Mn-$N^{\varepsilon 2}$(H26) | 2.17 | 2.10 | 2.13 | 2.14 | 2.10 | 2.07 | 2.07 | 2.07 | 2.26 | 2.10 |
| Mn-$N^{\varepsilon 2}$(H74) | 2.23 | 2.18 | 2.22 | 2.16 | 2.11 | 2.11 | 2.13 | 2.12 | 2.19 | 2.25 |
| Mn-$O^{\varepsilon 2}$(D159) | 2.13 | 2.03 | 2.11 | 2.11 | 2.15 | 2.16 | 1.95 | 1.94 | 2.44 | 2.15 |
| Mn-$N^{\varepsilon 2}$(H163) | 2.21 | 2.19 | 2.22 | 2.22 | 2.13 | 2.13 | 2.06 | 2.14 | 2.23 | 2.21 |
| Mn-O(WAT1) | - | 1.76 | 2.20 | - | 1.95 | 1.93 | 1.78 | 1.76 | 2.12 | 2.22 |
| Mn-$O^1$(LIG)[a] | 2.00 | - | - | - | - | - | - | - | - | - |
| Mn-$O^2$(LIG) | 2.34 | - | - | - | - | - | - | - | - | - |
| Mn-O(OL) | - | 1.80 | - | - | - | - | - | - | 1.82 | - |

| Neutron | Trp161Phe $D_2O_2$-Soaked[d] | | Trp161Phe $Mn^{3+}$SOD | | Trp161Phe $Mn^{2+}$SOD | |
|---|---|---|---|---|---|---|
| PDB ID | 8VHW | | 8VJ0 | | 8VHY | |
| Mn Bonds (Å) | A | B | A | B | A | B |
| Mn-$N^{\varepsilon 2}$(H26) | 2.13 | 2.20 | 2.01 | 2.01 | 2.20 | 2.11 |
| Mn-$N^{\varepsilon 2}$(H74) | 2.13 | 2.28 | 2.15 | 2.15 | 2.19 | 2.16 |
| Mn-$O^{\varepsilon 2}$(D159) | 2.20 | 2.05 | 2.01 | 2.01 | 2.13 | 2.20 |
| Mn-$N^{\varepsilon 2}$(H163) | 2.25 | 2.17 | 2.15 | 2.15 | 2.14 | 2.15 |
| Mn-O(WAT1) | 2.13 | - | 1.84 | 1.84 | 2.37 | 2.24 |
| Mn-$O^1$(LIG)[a] | - | 1.94 | - | - | - | - |
| Mn-$O^2$(LIG) | - | 2.61 | - | - | - | - |
| Mn-O(OL) | - | - | - | - | - | - |

[a]$O^1$(LIG) refers to the closest oxygen atom of the dioxygen species.

[b]Only chain A of $D_2O_2$-soaked Tyr34Phe MnSOD is bound by a dioxygen species, denoted as LIG. For chain A, an $^{-}$OD molecule (denoted OL for oxygen ligand) is observed binding opposite of Asp159 and is six-coordinate. Note that HERFD-XANES (**Supplementary Fig. 2a**) and EXAFS suggest a five-coordinate complex (**Supplementary Table 2**).

[c]For chain A of wildtype $Mn^{2+}$SOD, an $^{-}$OD molecule is observed binding opposite of Asp159 and is six-coordinate. Chain B is in the typical five-coordinated state. Note that HERFD-XANES suggest a five-coordinate complex (**Supplementary Fig. 2c**).

[d]Only chain B of $D_2O_2$-soaked Trp161Phe MnSOD is bound by a dioxygen species.

**Supplementary Table 4. Active site Mn bond lengths of DFT-optimized structures**

| Wildtype $Mn^{3+}$SOD (S=2) | | Tyr34Phe $Mn^{3+}$SOD (S=2) | |
|---|---|---|---|
| Bond | DFT (Å) | Bond | DFT (Å) |
| Mn-$N^{\varepsilon2}$(H26) | 2.06 | Mn-$N^{\varepsilon2}$(H26) | 2.04 |
| Mn-$N^{\varepsilon2}$(H74) | 2.13 | Mn-$N^{\varepsilon2}$(H74) | 2.11 |
| Mn-$N^{\varepsilon2}$(H163) | 2.06 | Mn-$N^{\varepsilon2}$(H163) | 2.09 |
| Mn-$O^{\delta2}$(D159) | 1.95 | Mn-$O^{\delta2}$(D159) | 1.98 |
| Mn-O(WAT1) | 1.80 | Mn-O(WAT1) | 1.82 |
| Wildtype $Mn^{2+}$SOD (S=5/2) | | Tyr34Phe $Mn^{2+}$SOD (S=5/2) | |
| Bond | DFT (Å) | Bond | DFT (Å) |
| Mn-$N^{\varepsilon2}$(H26) | 2.23 | Mn-$N^{\varepsilon2}$(H26) | 2.22 |
| Mn-$N^{\varepsilon2}$(H74) | 2.18 | Mn-$N^{\varepsilon2}$(H74) | 2.18 |
| Mn-$N^{\varepsilon2}$(H163) | 2.19 | Mn-$N^{\varepsilon2}$(H163) | 2.19 |
| Mn-$O^{\delta2}$(D159) | 2.06 | Mn-$O^{\delta2}$(D159) | 2.06 |
| Mn-O(WAT1) | 2.14 | Mn-O(WAT1) | 2.13 |
| Trp161Phe $Mn^{2+}$SOD + $HO_2^-$ (S=5/2) | | Tyr34Phe $Mn^{2+}$SOD + $HO_2^-$ (S=5/2) | |
| Bond | DFT (Å) | Bond | DFT (Å) |
| Mn-$N^{\varepsilon2}$(H26) | 2.24 | Mn-$N^{\varepsilon2}$(H26) | 2.28 |
| Mn-$N^{\varepsilon2}$(H74) | 2.16 | Mn-$N^{\varepsilon2}$(H74) | 2.20 |
| Mn-$N^{\varepsilon2}$(H163) | 2.15 | Mn-$N^{\varepsilon2}$(H163) | 2.16 |
| Mn-$O^{\delta2}$(D159) | 2.06 | Mn-$O^{\delta2}$(D159) | 2.17 |
| Mn-O(PEO) | 2.08 | Mn-O(PEO) | 2.10 |
| Tyr34Phe $Mn^{3+}$SOD + $HO_2^-$ (S=2) | | Tyr34Phe $Mn^{2+}$SOD + $HO_2^\bullet$ (S=3) | |
| Bond | DFT (Å) | Bond | DFT (Å) |
| Mn-$N^{\varepsilon2}$(H26) | 2.00 | Mn-$N^{\varepsilon2}$(H26) | 2.12 |
| Mn-$N^{\varepsilon2}$(H74) | 2.07 | Mn-$N^{\varepsilon2}$(H74) | 2.16 |
| Mn-$N^{\varepsilon2}$(H163) | 2.19 | Mn-$N^{\varepsilon2}$(H163) | 2.17 |
| Mn-$O^{\delta2}$(D159) | 2.01 | Mn-$O^{\delta2}$(D159) | 2.01 |
| Mn-O(PEO) | 1.82 | Mn-O(SUP) | 2.37 |

**Supplementary Table 5. Data collection and refinement statistics for Tyr34Phe MnSOD.**

| Data Collection Statistics | | | | | |
|---|---|---|---|---|---|
| | Neutron | | | X-ray | |
| Variant | Tyr34Phe MnSOD | | | | |
| Chemical State | $D_2O_2$-Soaked | Reduced | Oxidized | $H_2O_2$-Soaked | Reduced |
| PDB Code | 9BVY | 9BW2 | 9BWM | 9BWQ | 9BWR |
| Growth Conditions | Microgravity | | | Earth | |
| Diffraction Source | MaNDi | | | SSRL Beamline 14-1 | Rigaku FR-E SuperBright |
| Temperature (K) | 100 | 296 | 296 | 100 | 100 |
| Space group | $P6_122$ | | | $P6_122$ | |
| *a*, *b*, *c* (Å) | 78.69. 78.69, 239.94 | 79.4, 79.4, 238.43 | 79.29, 79.29, 240.61 | 78.22, 78.22, 239.50 | 78.09, 78.09, 235.19 |
| $\alpha, \beta, \gamma$ (°) | 90, 90, 120 | | | 90, 90, 120 | |
| Wavelengths (Å) | 2-4 | | | 0.9795 | 1.5418 |
| No. of images | 11 | 10 | 13 | 625 | 386 |
| Exposure time | 36 h | 28 h | 35 h | 1 s | 300 s |
| No. of unique reflections | 18698 | 14940 | 20703 | 85139 | 65175 |
| Total No. of reflections | 73477 | 123482 | 161187 | 170167 | 526729 |
| Resolution range (Å) | 14.84-2.30 (2.37-2.30) | 14.55-2.50 (2.58-2.5) | 14.39-2.28 (2.36-2.28) | 39.13-1.40 (1.45-1.40) | 50.00-1.50 (1.53-1.50) |
| Multiplicity | 4.1 (3.4) | 8.3 (6.7) | 7.79 (6.15) | 13.5 (13.6) | 8.1 (5.8) |
| I/σ(I) | 5.1 (2.7) | 6.6 (3.8) | 7.4 (3.7) | 7.3 (1.6) | 7.5 (1.38) |
| $R_{merge}$ | 0.277 (0.266) | 0.321 (0.292) | 0.246 (0.323) | - | - |
| $R_{meas}$ | 0.310 (0.305) | 0.340 (0.315) | 0.263 (0.349) | 0.039 (0.458) | 0.299 (0.816) |
| $CC_{1/2}$ | 0.783 (0.259) | 0.766 (0.117) | 0.953 (0.476) | 0.999 (0.788) | 0.847 (0.578) |
| $R_{pim}$ | 0.133 (0.142) | 0.106 (0.112) | 0.090 (0.127) | 0.027 (0.324) | 0.089 (0.411) |
| Completeness (%) | 90.4 (75.3) | 91.3 (90.7) | 97.78 (88.40) | 98.8 (97.5) | 94.4 (85.2) |
| Refinement Statistics | | | | | |
| $R_{work}$ | 0.265 | 0.293 | 0.242 | 0.184 | 0.189 |
| $R_{free}$ | 0.299 | 0.330 | 0.275 | 0.205 | 0.210 |
| [a]No. of atoms | | | | | |
| Protein | 6547 | 6233 | 6617 | 3160 | 3160 |
| [b]Solvent | 1068 | 496 | 147 | 571 | 782 |
| Mn | 2 | 2 | 2 | 2 | 2 |
| R.m.s. deviations | | | | | |
| Bond lengths (Å) | 0.002 | 0.006 | 0.001 | 0.004 | 0.003 |
| Bond angles (°) | 0.541 | 0.522 | 0.505 | 0.683 | 0.638 |
| Average *B*-factor | 3.69 | 15.36 | 35.45 | 19.65 | 18.38 |
| Protein | 3.73 | 15.45 | 35.42 | 18.42 | 15.41 |
| Water | 3.45 | 13.87 | 36.74 | 25.90 | 30.07 |
| Mn | 2.38 | 14.70 | 42.91 | 14.21 | 9.99 |
| Peroxide | 2.98 | - | - | 16.82 | - |
| [c]Coordinate Error (Å) | 0.30 | 0.42 | 0.30 | 0.15 | 0.15 |

[a]No. of atoms is inclusive of H/D atoms for neutron structures.

[b]The neutron structure of reduced and oxidized Tyr34Phe were collected at room temperature and, as a result, have fewer solvent atoms compared to the other structures.

[c]Estimated coordinate error determined by PHENIX. Note that the disorder of atom positions is reflected by the *B*-factor.
